# Novel epidemiological model of gastrointestinal-nematode infection to assess grazing cattle resilience by integrating host growth, parasite, grass and environmental dynamics

**DOI:** 10.1101/2022.05.14.491963

**Authors:** J.A.N. Filipe, I. Kyriazakis, C. McFarland, E.R. Morgan

**Affiliations:** Biomathematics & Statistics Scotland, Rowett Institute of Nutrition and Health, University of Aberdeen, AB25 2ZD, UK; Institute for Global Food Security, Queen’s University Belfast, Biological Sciences, 19, Chlorine Gardens, BT9 5DL, UK

**Keywords:** helminth, *Ostertagia ostertagi*, *Cooperia oncophora*, climate, parasite-induced anorexia, mathematical model

## Abstract

Gastrointestinal nematode (GIN) infections are ubiquitous and often cause morbidity and reduced performance in livestock. Emerging anthelmintic resistance and increasing change in climate patterns require evaluation of alternatives to traditional treatment and management practices. Mathematical models of parasite transmission between hosts and the environment have contributed towards the design of appropriate control strategies in ruminants, but have yet to account for relationships between climate, infection pressure, immunity, resources, and growth. Here, we develop a new epidemiological model of GIN transmission in a herd of grazing cattle, including host tolerance (body weight and feed intake), parasite burden and acquisition of immunity, together with weather-dependent development of parasite free-living stages, and the influence of grass availability on parasite transmission. Dynamic host, parasite and environmental factors drive a variable rate of transmission. Using literature sources, the model was parametrised for *Ostertagia ostertagi*, the prevailing pathogenic GIN in grazing cattle populations in temperate climates. Model outputs were validated on published empirical studies from first season grazing cattle in Northern Europe. These results show satisfactory qualitative and quantitative performance of the model; they also indicate the model may approximate the dynamics of grazing systems under co-infection by *O. ostertagi* and *Cooperia oncophora*, a second GIN species common in cattle. In addition, model behaviour was explored under illustrative anthelmintic treatment strategies, considering impacts on parasitological and performance variables. The model has potential for extension to explore altered infection dynamics as a result of management and climate change, and to optimise treatment strategies accordingly. As the first mechanistic model to combine parasitic and free-living stages of GIN with host feed-intake and growth, it is well suited to predict complex system responses under non-stationary conditions. We discuss the implications, limitations and extensions of the model, and its potential to assist in the development of sustainable parasite control strategies.

**HIGHLIGHTS:**

- Nematode control in cattle is complicated by drug resistance and climate change
- A model was developed to predict GIN epidemiology under varying conditions
- The model incorporates cattle growth, infection and immunity, grass availability, weather
- Predictions were validated against empirical studies of GIN in N Europe
- The model applies to *Ostertagia ostertagi*, and possibly to co-infecting *Cooperia*

https://www.lmcni.com/

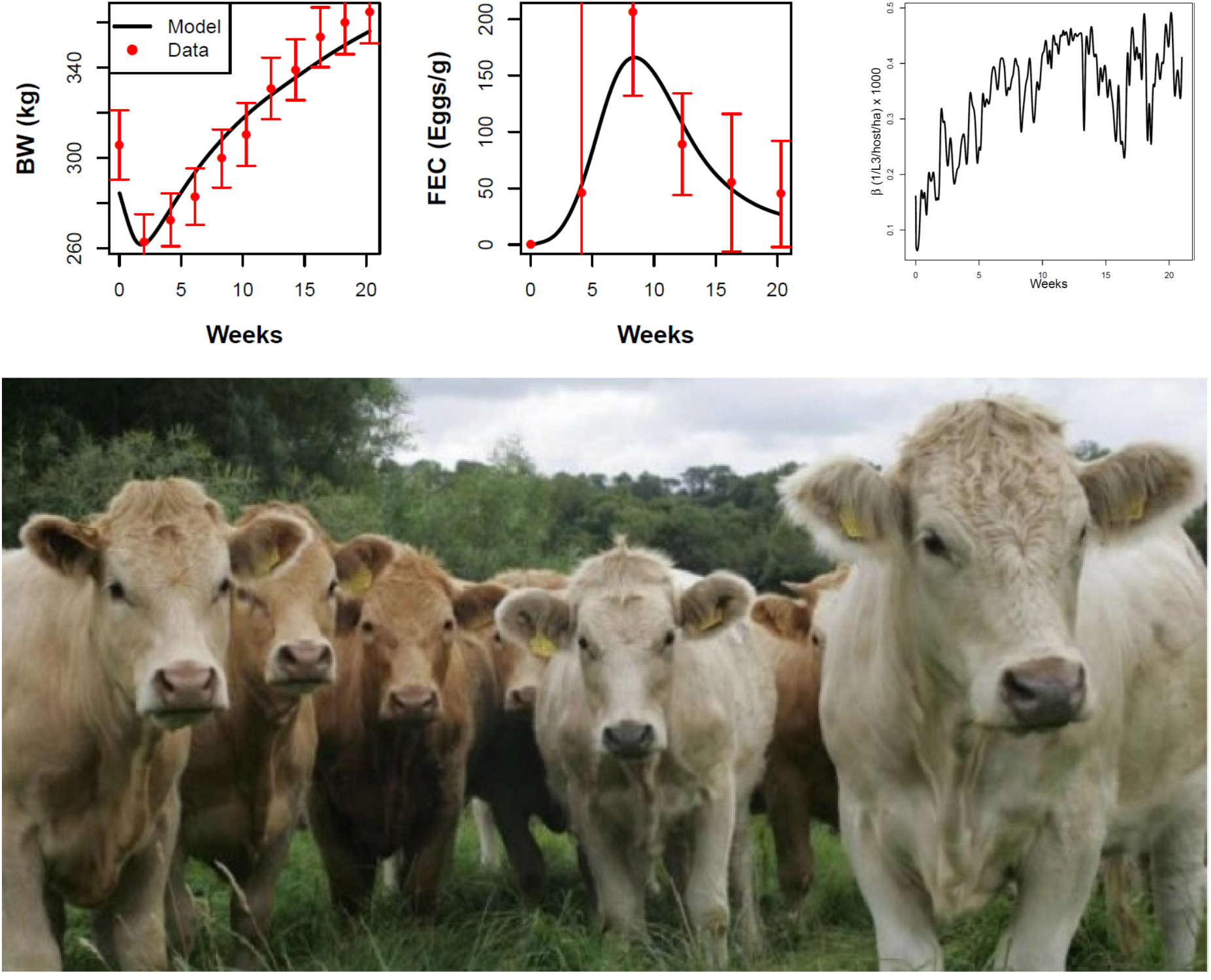

## 1. Introduction

Gastrointestinal nematode (GIN) infections have significant health, welfare and economic impacts in cattle and other grazing-livestock species, often through the occurrence of sub-clinical disease (Armour, 1980; Charlier et al., 2020b; Fox, 1993). The dominant GINs of cattle in temperate climates are *O. ostertagi* and *Cooperia oncophora*, which cause parasitic gastroenteritis primarily in first grazing season (FGS) cattle (Forbes, 2020; Michel, 1969). The use of anthelmintic treatments remains a first line practice to safeguard the health and growth performance of grazing livestock. However, the sustainability of this practice is threatened by the emergence of anthelmintic resistance in cattle worldwide (Kaplan and Vidyashankar, 2012; Rose Vineer et al., 2020a) requiring alternative parasite control strategies to limit further development of resistance. Further challenges emerge with the increasing pace of climate change, which may affect parasite development, grass availability and host growth; these challenges require further decisions on how to adapt the management practices of grazing livestock (Skuce et al., 2013; Vercruysse et al., 2018). Mathematical models are an important tool for evaluating and comparing alternative treatment and management strategies given the practical difficulties of doing so in experiments (Smith, 2011). Such models are useful, for example, for evaluating targeted selective treatments applied to individuals on the basis of parasitological or performance indicators (Charlier et al., 2014; Höglund et al., 2013); and where the benefits of preserving parasite refugia (Hodgkinson et al., 2019; van Wyk, 2001) are balanced against the risks of reduced health and performance in untreated animals. Achieving these goals requires the availability of models that include the full life cycle of the parasite, as well as the dynamics of immunity, grass availability and consumption, and animal growth.

Models of the full-cycle transmission of the most pathogenic GIN in cattle, *Ostertagia ostertagi*, have been developed and applied to field data (Grenfell et al., 1987a; Smith and Grenfell, 1994), but while these models have incorporated host and free-living (FL) parasite stages and host acquired immunity, they have not included host performance traits (Smith, 1997). However, weight gain and feed intake are important variables in the host-parasite interaction as well as having economic significance due to reduction in gain, especially in young parasitized cattle (Bell et al., 1988; Coop and Kyriazakis, 1999; Symons, 1985). In practice, body weight, as well as parasite eggs in host faeces, can be monitored during grazing to guide the applications of anthelmintic treatment; and affordable technology for routine individual weighing is becoming increasingly available (González-García et al., 2018). Weight and intake have important roles in system behaviour and models thereof, not only as output variables but also because they affect parasite epidemiology. First, the rate at which infective larvae are ingested (transmission rate) (Grenfell, 1988) is controlled by the rate of feed intake, of which weight is a main determinant (NRC, 1987); it is also controlled by the density of grass on pasture (Henriksen et al., 1976; Nansen et al., 1988). Second, intake is reduced through parasite-induced anorexia (Bell et al., 1988; Coop and Kyriazakis, 1999). Models that incorporate host performance during infection with *O. ostertagi* have been developed (Berk et al., 2016a), but have not yet been incorporated with a realistic and parameterised model of the parasite FL stages, which has been developed separately (Rose et al., 2015). The aim of our paper is to contribute to the above goals by integrating these system layers, whilst also aiming to focus on fewer host performance variables than the previous models, in the interests of transparent model behaviour and simpler parameterisation.

Here, building on elements from the above models, we propose a dynamic transmission model of the full parasite lifecycle, parameterised for *O. ostertagi* in cattle using parameter estimates from literature sources. The model incorporates 1) parasite load, acquired immunity, and weight and feed intake as host variables, 2) FL parasite stages influenced by local weather and climate, and 3) variable grass biomass. This model allows, for the first time, to investigate the consequences of control practices on both parasitological and performance variables, while taking into account of variability in weather and seasonality in climate. We tested (validated) model predictions against field data from several studies in Northern Europe; these studies took place during the FGS and under natural infection and immunity progression. In addition, we explored the potential of the model to predict the impacts of simplified anthelmintic treatments, leaving the effects of alternative anthelmintic treatments for later consideration. We anticipate there is potential to parameterise the model for other GIN species of grazing ruminants, and to explore behaviour under future climate change. We hypothesise that interactions between growth, grass availability and intake, infection and immunity, and the dynamics of the parasite pasture stages, lead to interpretable non-linear responses in system behaviour, and that these can be explored to enhance the outcomes of treatment interventions. Further, we hypothesise that when calibrated to conditions in published experimental trials in FSG cattle, the model will reproduce observed patterns of animal infection and performance.

## 2. Materials and Methods

### 2.1 Overview of the full transmission model

The model of GIN transmission in a herd of grazing cattle, including the full lifecycle of the parasite, links four sub-models schematised in Fig. 1. Two sub-models describe the animal host and its interaction with the parasite. First, the host infection and immunity sub-model (Section 2.2) describes host daily ingestion of infective third stage larvae (L3) on herbage, L_3h_, from the current grass G on pasture (determined by the daily grass intake, i.e. feed intake, FI, and its contamination L_3h_/G), the development of parasitic larval stages L (including L4 and L5) into adult worms W that produce eggs excreted via the host faeces onto pasture, Ep, and the development of acquired immunity, I_m_, by the host concomitantly with its gradual infection. Second, the host growth sub-model (Section 2.3) describes the bodyweight (BW), FI and growth of the host given its genetic propensity for growth, age, current level of infection (since the start of grazing, or turnout onto pasture), and the current immune state and response to the parasite loads L and W. The daily grass intake FI is calculated based on the current BW and maintenance functions; the faecal mass output is the non-digested intake and is used to calculate the current number of parasite eggs excreted per unit mass (faecal egg counts) given the current number of eggs produced by the resident worms.

**Fig. 1.**
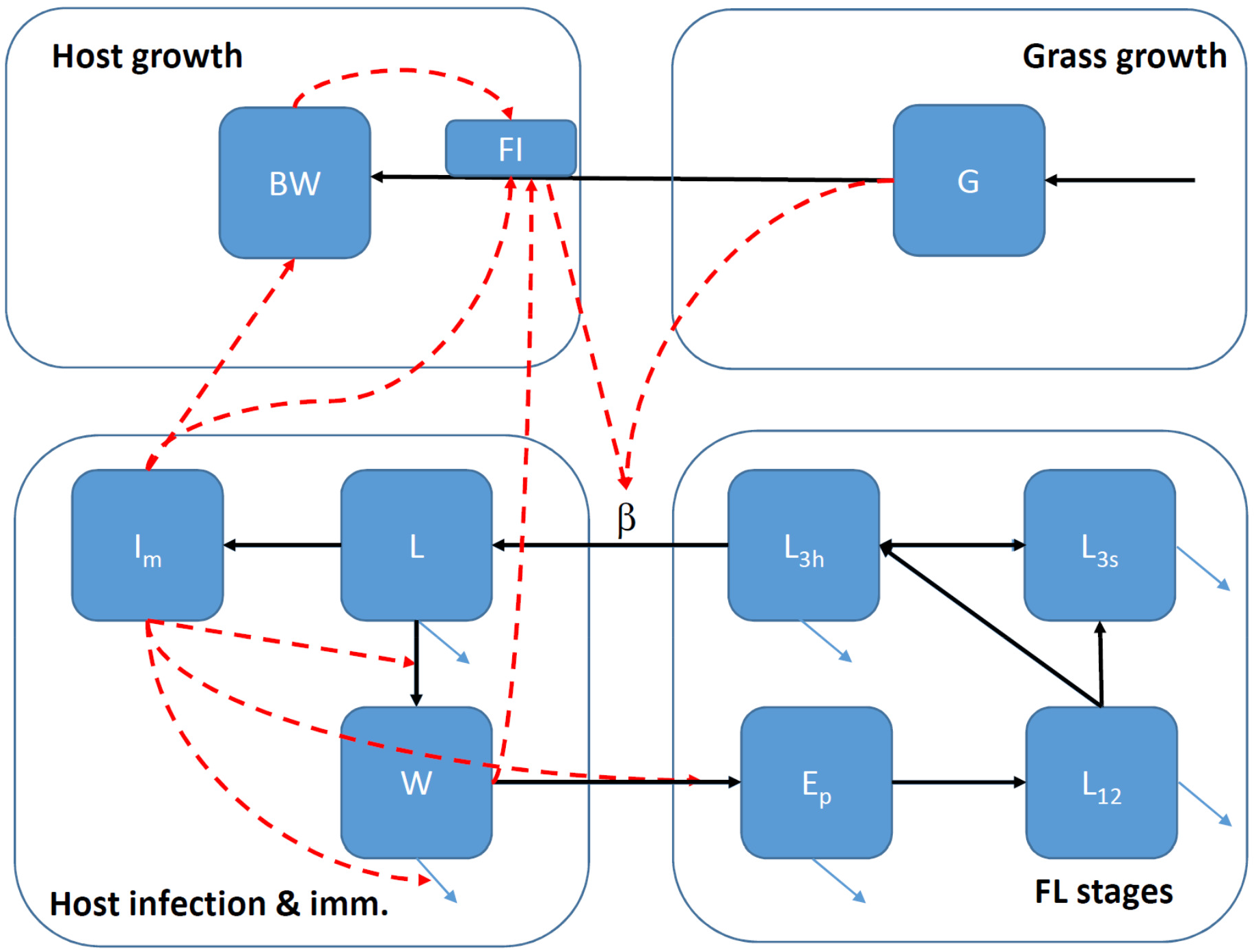
Structure of the full-cycle model of GIN transmission. The model comprises four sub-models representing: host growth; grass growth; host infection and the development of immunity; and the free living (FL) parasite stages. The details of each sub-model are given in the text, Sections 2.2 to 2.5. Squares: state variables. Arrows: flow or transition (black), mortality (blue), influence (red).

Two further sub-models describe the grazing environment and the survival of the FL parasite population. First, the grass growth model (Section 2.4) describes the availability of grass G through the grazing season in terms of dry mass per unit area of pasture, balancing grass growth and consumption through gazing by the herd; a net decrease in G will increase the larval concentration on herbage, L_3h_, and its ingestion. Increases in intake during the host’s growth trajectory, as well as parasite-induced anorexia will also affect L3 ingestion. Second, the FL-stages model (Section 2.5) describes the development of parasite stages outside the host, from eggs excreted in host faeces, through intermediate first and second larval stages, L12, that reside within the faeces, and into infective larvae on pasture, which migrate in both directions between herbage, L_3h_, and soil, L3s, and while present on grass can be ingested through grazing. The development, survival and migration of these life stages depends on daily temperature and rainfall. The dynamics of the full transmission cycle are summarised by the parasite’s effective reproduction number, R_e_, which incorporates magnification or decline from each lifecycle stage through the effects of the weather and host-parasite interaction.

The full transmission model was built by linking the four sub-models and establishing suitable interactions between host, parasite and grass variables, and between the hosts in the herd who share the FL parasites and the available grass. For tractability, the implementation of the model is deterministic, i.e. each set of input conditions generates a single model prediction. For simplicity, the grazing host population (herd) is characterised by a stocking density per hectare, and the host sub-models are assumed to represent the average state of the animals in the herd; individual demographic stochasticity is not included. The grazing movement and the grass intake and L3 ingestion by the animals are represented in an average sense, assuming spatially-mixed grass consumption and faecal deposition across the pasture; this is a mean-field rather than a spatially-explicit representation of the host-environment interactions. Parasites in all stages are also treated at a population mean level. These assumptions are shared by the past mathematical models of GINs in FGS cattle that we have built on, which have similarly focused on *O. ostertagi* (Berk et al., 2016a; Grenfell et al., 1987a; Rose et al., 2015). Each sub-model and supporting literature are described in detail next. All state variables and parameters of the model are described in Tables 1-4.

**Table 1.** Host infection and immunity. State variables and parameters defined in sub-model 1 (Section 2.2). State variables are time dependent. Parameters are constant or a function of the immunity level.

| Variable | Description | Units | Comments |  |
| --- | --- | --- | --- | --- |
| J | intake of L3 larvae | larvae/d | Eq. (1) |  |
| $L_{3c}$ | L3 concentration on grass | larvae/kgDM | $L_{3c}=L_{3h}/(G/A_g)$ sub-model 3 | |
| $L_{3h}$ | L3 density on herbage | larvae/ha | Sub-model 4 | |
| FIDM | Feed intake dry matter (DM) | kg DM/d | Sub-model 2 |  |
| $L_i$ | Development compartments between L3 and L | - | Number within host, $i=1\dots n_L$ | |
| L | Larvae stage pre-adult (fourth and fifth) | - | Number within host ($L=L_{nL}$) | |
| W | Adult worms in host | - | - |  |
| E | Cumulative number of eggs produced by adults | - | - |  |
| Epd | Eggs produced daily by female worms | eggs/d | - |  |
| FEC | Average faecal egg counts | eggs/g | per daily wet faeces (Eq. (6)) |  |
| C | Marker of cumulative exposure to L3 | - | Unbounded |  |
| $I_m$ | Level of acquired immunity of the host | - | Bounded between 0 and 1 | |
| Parameter |  | Units | Value | Source |
| $\varepsilon(I_m)$ | Establishment probability from L3 to adult | - | - | <sup>1</sup>, Eq. (7) |
| $\varepsilon'$ | Per-compartment establishment probability of L3 | - | $\varepsilon' = \varepsilon^{1/n_L}$ | - |
| $\varepsilon_0$ | Establishment probability of L3 (naïve animal) | - | 0.60 | <sup>2</sup> |
| $\varepsilon_1$ | Establishment probability of L3 (immune host) | - | 0.05 | <sup>2</sup> |
| $\sigma$ | Development rate of L3 | 1/d | $n_L/T_L$ | - |
| $T_L$ | Mean pre-patency | d | 32 | <sup>3</sup> |
| $n_L$ | Number of L<sub>i</sub> development compartments | - | 5 | This paper |
| $\mu(l_m)$ | Mortality rate of adult worms | 1/d | - | Eq. (7) |
| $\mu_0$ | Mortality rate of adult worms (naïve host) | worms/d | 0.025 | <sup>2, 4, 5</sup> |
| $\mu_1$ | Mortality rate of adult worms (immune host) | larvae/d | 0.06 | <sup>2, 6</sup> |
| $p_f$ | Proportion of female adult worms | - | 0.55 | <sup>2</sup> |
| $f_e(l_m, W)$ | Effective fecundity rate of female adults worms | eggs/d | - | <sup>7, 8</sup> |
| $W_s$ | Adult worm scale of density dependence | - | 15000 | <sup>7, 9</sup> |
| $f(l_m)$ | Fecundity rate of worms (no density dependence) | eggs/d | - | Eq. (7) |
| $f_0$ | Fecundity rate of worms (naïve host) | eggs/d | 350 | <sup>7, 10</sup> |
| $f_1$ | Fecundity rate of worms (immune host) | eggs/d | 30 | <sup>7, 10</sup> |
| $n_f$ | Exponent relating f to $l_m$ | - | 0.2 | This paper |
| $l_m(0)$ | Initial level of immunity of host | - | 0 | FGS, or as stated |
| $C_m$ | Cum. L3 exposure at ~25% of maximum immunity | - | 70k | <sup>2, 6</sup> |
| $\sigma_C$ | Development rate of C | 1/d | $n_C/T_C$ | - |
| $T_C$ | Mean time of development of C | d | 32 | See $T_L$ |
| $n_C$ | Number of C development compartments | - | 5 | This paper |
| $\mu_C$ | Rate of loss of host immunity | 1/d | $\ln(3)/180$ | <sup>11</sup> |

<sup>1</sup> Includes mortality or arrest at rate $(1-\epsilon')$ . $\sigma$ . Rate of transition to the next compartment: $\epsilon'$ . $\sigma$ .
<sup>2</sup> Verschave et al. (2014)
<sup>3</sup> Assuming the observed 21d (Anderson, 2000; Verschave et al., 2014) corresponds to the 25<sup>th</sup> percentile of the gamma distributed development time assumed in Eq. (9) with shape parameter $n_C$ .
<sup>4</sup> Michel et al. (1973)
<sup>5</sup> Grenfell et al. (1987b)
<sup>6</sup> Smith (1994)
<sup>7</sup> Berk et al. (2016a)
<sup>8</sup> Bishop and Stear (1997)
<sup>9</sup> Michel (1969)
<sup>10</sup> Smith et al. (1987)
<sup>11</sup> Mapped from an assumption of 70% decay in overall immunity level in 180d (Rose Vineer et al., 2020b).

**Table 2.**
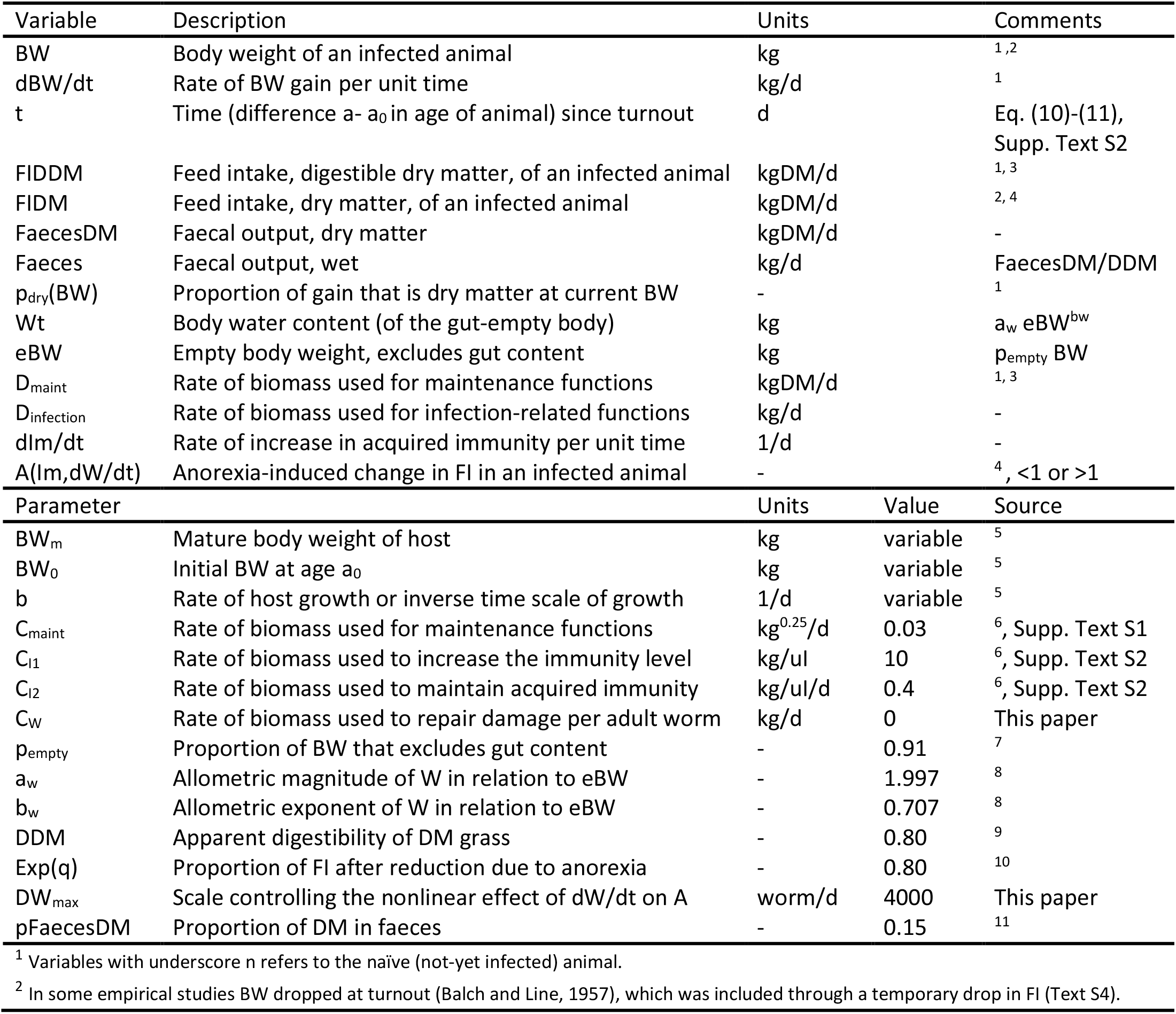
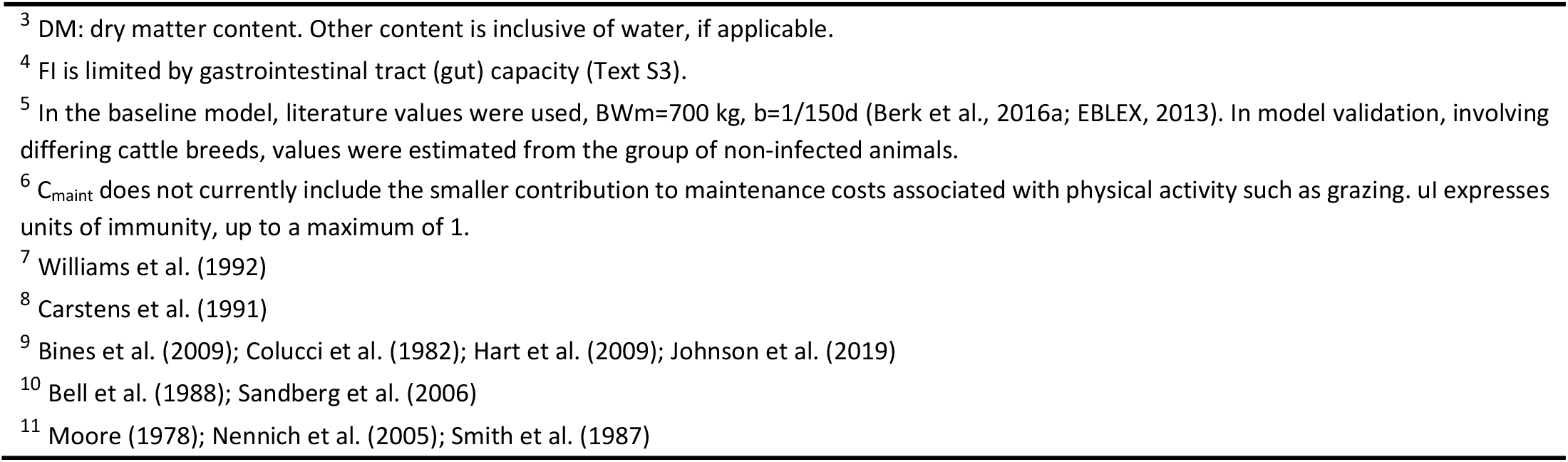
Host growth. State variables and parameters defined in sub-model 2 (Section 2.3). State variables are time dependent or may depend explicitly on other variables, e.g. p_dry_(BW).

| Variable | Description | Units | Comments |  |
| --- | --- | --- | --- | --- |
| BW | Body weight of an infected animal | kg | 1, 2 |  |
| $\text{dBW}/\text{dt}$ | Rate of BW gain per unit time | kg/d | 1 | |
| $t$ | Time (difference $a - a_0$ in age of animal) since turnout | d | Eq. (10)-(11),<br>Supp. Text S2 | |
| FIDDM | Feed intake, digestible dry matter, of an infected animal | kgDM/d | 1, 3 |  |
| FIDM | Feed intake, dry matter, of an infected animal | kgDM/d | 2, 4 |  |
| FaecesDM | Faecal output, dry matter | kgDM/d | - |  |
| Faeces | Faecal output, wet | kg/d | FaecesDM/DDM |  |
| $p_{\text{dry}}(\text{BW})$ | Proportion of gain that is dry matter at current BW | - | 1 | |
| $W_t$ | Body water content (of the gut-empty body) | kg | $a_w e\text{BW}^{b_w}$ | |
| eBW | Empty body weight, excludes gut content | kg | $p_{\text{empty}} \text{BW}$ | |
| $D_{\text{maint}}$ | Rate of biomass used for maintenance functions | kgDM/d | 1, 3 | |
| $D_{\text{infection}}$ | Rate of biomass used for infection-related functions | kg/d | - | |
| $\text{dIm}/\text{dt}$ | Rate of increase in acquired immunity per unit time | 1/d | - | |
| $A(\text{Im}, \text{dW}/\text{dt})$ | Anorexia-induced change in FI in an infected animal | - | 4, <1 or >1 | |
| Parameter |  | Units | Value | Source |
| $\text{BW}_m$ | Mature body weight of host | kg | variable | 5 |
| $\text{BW}_0$ | Initial BW at age $a_0$ | kg | variable | 5 |
| $b$ | Rate of host growth or inverse time scale of growth | 1/d | variable | 5 |
| $C_{\text{maint}}$ | Rate of biomass used for maintenance functions | $\text{kg}^{0.25}/\text{d}$ | 0.03 | 6, Supp. Text S1 |
| $C_{I1}$ | Rate of biomass used to increase the immunity level | kg/ul | 10 | 6, Supp. Text S2 |
| $C_{I2}$ | Rate of biomass used to maintain acquired immunity | kg/ul/d | 0.4 | 6, Supp. Text S2 |
| $C_w$ | Rate of biomass used to repair damage per adult worm | kg/d | 0 | This paper |
| $p_{\text{empty}}$ | Proportion of BW that excludes gut content | - | 0.91 | 7 |
| $a_w$ | Allometric magnitude of $W$ in relation to eBW | - | 1.997 | 8 |
| $b_w$ | Allometric exponent of $W$ in relation to eBW | - | 0.707 | 8 |
| DDM | Apparent digestibility of DM grass | - | 0.80 | 9 |
| $\text{Exp}(q)$ | Proportion of FI after reduction due to anorexia | - | 0.80 | 10 |
| $\text{DW}_{\text{max}}$ | Scale controlling the nonlinear effect of $\text{dW}/\text{dt}$ on $A$ | worm/d | 4000 | This paper |
| $p_{\text{FaecesDM}}$ | Proportion of DM in faeces | - | 0.15 | 11 |

<sup>1</sup> Variables with underscore n refers to the naïve (not-yet infected) animal.
<sup>2</sup> In some empirical studies BW dropped at turnout (Balch and Line, 1957), which was included through a temporary drop in FI (Text S4).
<sup>3</sup> DM: dry matter content. Other content is inclusive of water, if applicable.
<sup>4</sup> FI is limited by gastrointestinal tract (gut) capacity (Text S3).
<sup>5</sup> In the baseline model, literature values were used, BWm=700 kg, b=1/150d (Berk et al., 2016a; EBLEX, 2013). In model validation, involving differing cattle breeds, values were estimated from the group of non-infected animals.
<sup>6</sup> $C_{\text{maint}}$ does not currently include the smaller contribution to maintenance costs associated with physical activity such as grazing. $u_I$ expresses units of immunity, up to a maximum of 1.
<sup>7</sup> Williams et al. (1992)
<sup>8</sup> Carstens et al. (1991)
<sup>9</sup> Bines et al. (2009); Colucci et al. (1982); Hart et al. (2009); Johnson et al. (2019)
<sup>10</sup> Bell et al. (1988); Sandberg et al. (2006)
<sup>11</sup> Moore (1978); Nennich et al. (2005); Smith et al. (1987)

**Table 3.**
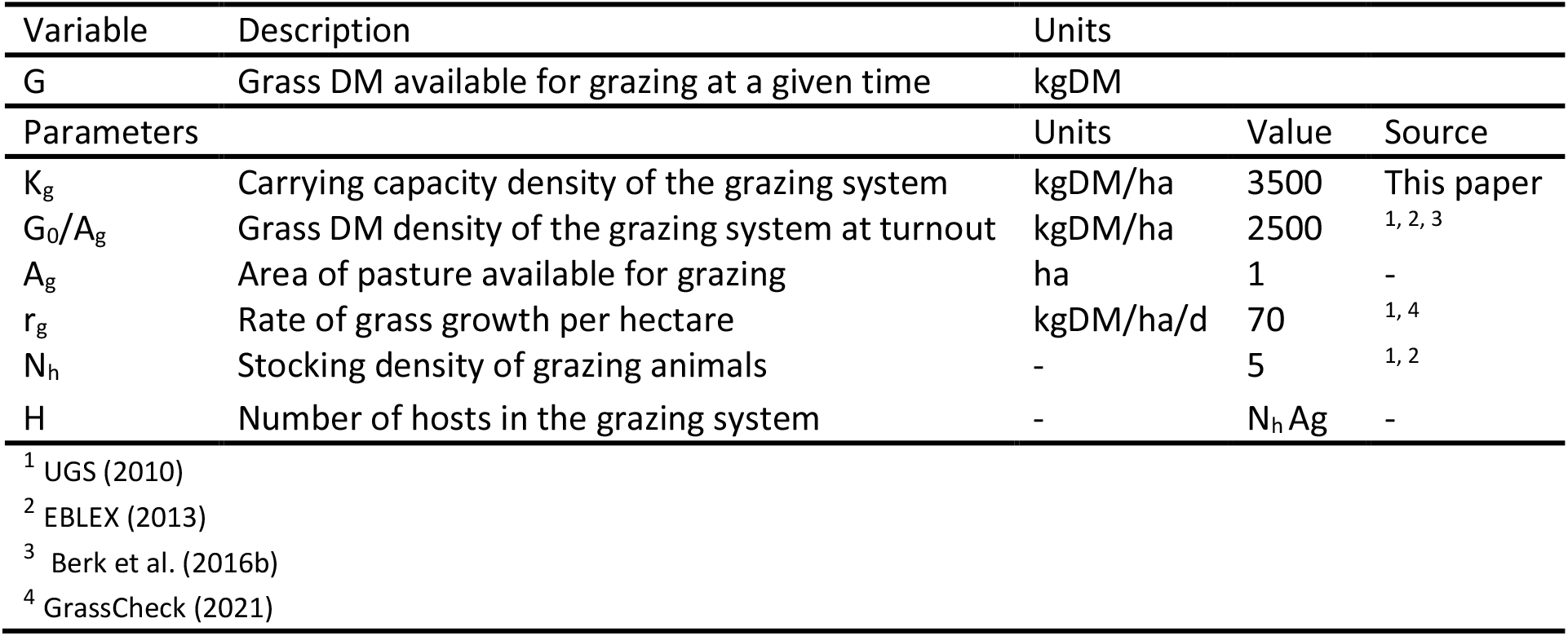
Grass growth. State variables and parameters defined in sub-model 3 (Section 2.4).

**Table 4.** Free-living stages. State variables and parameters defined in sub-model 4 (Section 2.5) by Rose et al. (2015). The dependency of the parameters on temperature and precipitation is given in Supplementary Table S1.

| Variable | Description | Units | Comment |
| --- | --- | --- | --- |
| $E_p$ | Density of eggs on pasture | eggs/ha | - |
| $E_c$ | Cumulative eggs deposited on pasture | eggs/ha | - |
| $L_{12}$ | Density of L1 and L2 larvae in faeces on pasture | larvae/ha | - |
| $L_{3f}$ | Density of L3 in faeces on pasture | larvae/ha | - |
| $L_{3p}$ | Density of L3 in pasture migrated from faeces | larvae /ha | herbage and soil |
| $L_{3h}$ | Density of L3 on herbage on pasture | larvae /ha | As in Table 1 |
| $L_{3s}$ | Density of L3 in soil | larvae /ha | - |
| Parameters |  | Units | Weather dependent |
| $\beta(\tau)$ | Probability of ingestion (transmission) per L3 per host | 1/d | Yes, new variable |
| $\delta$ | Rate of development from egg to L3 | eggs/d | Yes (Supp. Table S1) |
| $\mu_1$ | Rate of mortality of eggs on pasture | eggs/d | *idem* |
| $\mu_2$ | Rate of mortality of L1 and L2 larvae in faeces | larvae/d | *idem* |
| $\mu_3$ | Rate of mortality of L3 larvae in faeces | larvae/d | *idem* |
| $\mu_4$ | Rate of mortality of L3 larvae in soil | larvae/d | *idem* |
| $\mu_5$ | Rate of mortality of L3 larvae on herbage | larvae/d | *idem* |
| $m_1$ | Rate of herbage-soil migration of L3 | larvae/d | *idem* |
| $m_2$ | Proportion of pasture L3 that are on herbage | - | *idem* |

### 2.2 Sub-model 1: Host infection and immunity

The model dynamics of host infection are as follows. The influx (J) of third stage infective larvae (L3) by a grazing animal at a given time (t) during the FGS is given by

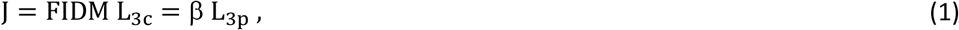

where FIDM is the animal’s rate of dry matter intake and L3c is the concentration of L3 on grass (as distinct from the density of L3 per unit area, L_3h_; Table 1). The second equality in Eq. (1) is for later use; it involves the rate of transmission per infective larva (β) and the density of L3 on pasture (L_3p_). In cases where a dose of L3 is inoculated at turnout, as part of experimental trials used to validate the model, there is an additional pulse in J at t=0. Ingested L3 that survive during establishment develop into stage L (combined stages L4 and L5) before developing to dioecious adult worms. This development is represented through nL mathematical compartments, or phases (L_i_, i=1…n_L_), that confer a gamma distribution to its time duration:

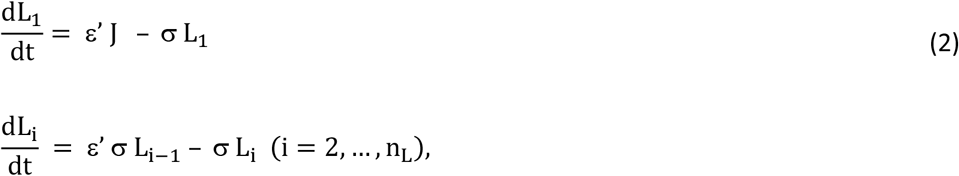

with L= L_nL_ and ε’ the probability of establishment (Table 1). This gamma distribution (in fact Erlang) is used instead of the common exponential distribution (n_L_=1) in order to ensure that pre-patency does not end prematurely and has the expected duration (Leclerc et al., 2014). The choice nL=5 ensures also that the distribution of times is approximately normal. Stages L become adult worms (W) at rate

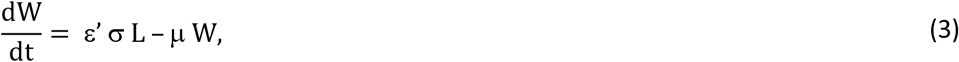

Eggs are produced by female adult worms at rate (Epd)

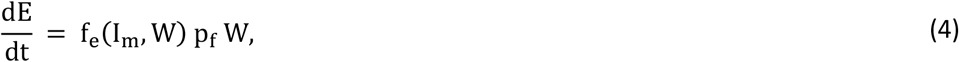

where f_e_ is the effective fecundity rate, which is reduced by the level of acquired immunity (I_m_) and by the worm density

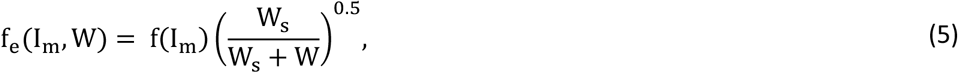

and where fecundity f(I_m_) is constrained by immunity but not by density. We modified the form of this density dependence in relation to (Berk et al., 2016a; Bishop and Stear, 1997) such that f_e_ = f(I_m_) when W is small.

The faecal egg count (FEC) is the egg output per gram of daily wet faecal output

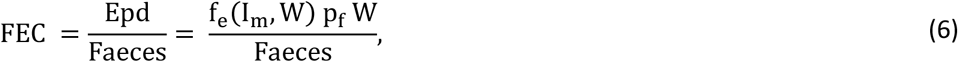

where Faeces (Table 2) is the wet faecal output (calculated in Eq. 22) expressed as grams/d. FEC observations are usually based on faecal samples across the herd or pasture and thus represent an average over the grazing herd.

Three within-host parasite traits, L3 establishment, adult worm mortality, and female worm fecundity, are regulated by the host’s immune response (Churcher et al., 2006; Grenfell et al., 1987b; Smith et al., 1987), each of which shifts between two parasite-specific limits as the level of immunity increases (Berk et al., 2016a):

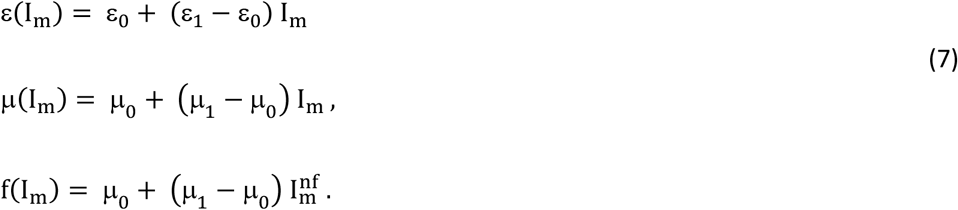

We assume that the first two responses develop at equal speed and that the reduction in worm fecundity occurs faster, via the exponent n_f_ <1 suggested by empirical observations (Dorny et al., 1997; Smith et al., 1987).

The level of acquired immunity is assumed to be bounded between 0 and 1; it is given by a sigmoidal growth function (here a von Bertalanffy-type function) of the cumulative exposure to L3 (C), given by

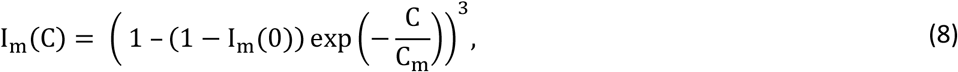

where I_m_(0) is the level of immunity at turnout, from when cumulative exposure is measured. In the FGS we expect I_m_(0)=0.

The cumulative exposure C is a hypothetical memory of antigen stimulation from the incrementally ingested L3 (Smith and Grenfell, 1985); it emerges after a time delay required for the development of acquired effector mechanisms. The dynamics of C are represented in a similar way to the development of the within-host parasite stages,

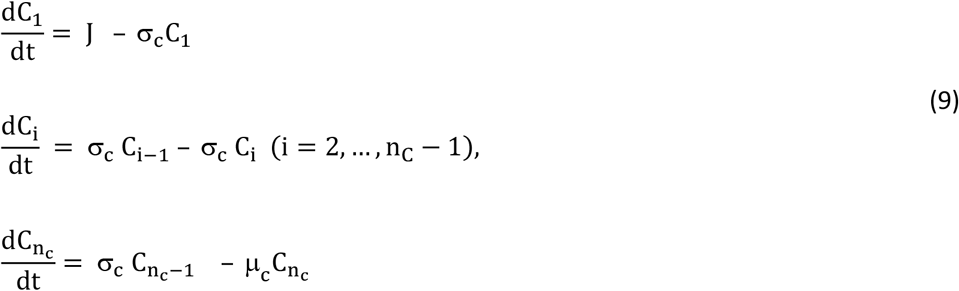

with C=C_nc_, and where we allow for loss of immunity through a constant-rate loss in C (μ_c_). The equations for C are similar to those for L, but without mortality terms and with distinct rate parameters (Table 1). This representation builds on previous work (Anderson and May, 1985; Roberts and Grenfell, 1991; Smith and Grenfell, 1994) that related the theoretical level of acquired immunity directly to the cumulative number of L3 ingested and where immunity had a constant rate of loss. We expressed loss of immunity similarly, through a loss in the cumulative marker C; but rather than using C directly as immunity level, we used the bounded immunity level I_m_ (Eq. 8) for ease of interpretation because a given level of C is not directly interpretable (Rose Vineer et al., 2020b). A second difference in our representation is the distribution of the temporal delay in the emergence of C upon exposure; Eq. (9) prevents premature emergence by disallowing a skew towards zero. In addition, we assumed that the time scale of development of immunity is the same as that of the development of L3 into adult worms (Table 1); this is a minimal working assumption as immunity could develop faster or more slowly than the with-host parasite stages; however, we have no evidence in favour of either case. A similar remark applies to the time delay; amid lack of evidence we assumed that the number of development compartments nC is the same as nL in the development of adult worms, which could be revised amid fresh evidence. Hypobiotic arrest and re-emergence of parasitic larvae, which is affected by season and immunity (Charlier et al., 2020a) was not included since this comes into play only towards the end of the grazing season, and is in any case too poorly understood to parameterise.

### 2.3 Sub-model 2: Host growth

The variables and parameters of this sub-model are detailed in Table 2. The body weight (BW_n_) of a naïve host (FGS calf that has not yet been infected) is assumed to increase with age according to a Gompertz function (Berk et al., 2016a; Forni et al., 2009),

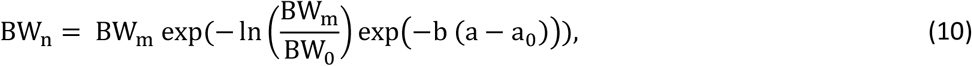

where the growth rate (b) and mature weight (BW_m_) are performance parameters inherent to the host species and breed, and BW_0_ and a_0_ are weight and age at turnout. The rate of weight gain is given by

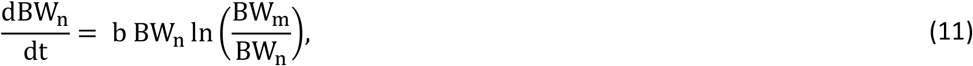

where t is time since turnout. The daily digestible (D) DM feed intake (FIDDM n) is utilised by the animal as DM weight gain and in dry-mass flows associated with maintenance functions (D_maint,n_):

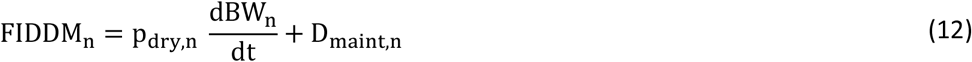

where p_dry,n_ is the proportion of gain that is DM, and D_maint,n_ is expressed as DM.

An infected animal has reduced growth in relation to its potential growth if uninfected (Eq. (10)) because it has reduced intake (parasite-induced anorexia) and there are costs associated with infection (Coop and Kyriazakis, 1999; Coop et al., 1977; Fox et al., 1989). To derive the body weight (BW) and feed intake (FIDDM) of the infected animal we assume that, at a given time, its intake is suppressed by a factor A<1 in relation to that of the naïve (uninfected) animal of the same weight,

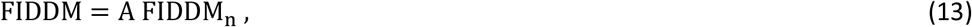

where 1-A is the proportion of reduction in FI caused by anorexia; it is also possible to have A>1 during compensatory growth. We assume, furthermore, that intake resources are used in additional mass flows (Coop and Kyriazakis, 1999) associated with functions that tackle infection (D_infection_):

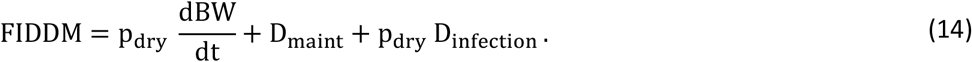

We expect the maintenance costs of the infected and naïve animals to be the same when they have the same BW. Rewriting Eq. 14 and substituting in Eqs. 11-13, we derive the rate of gain of the infected animal as being:

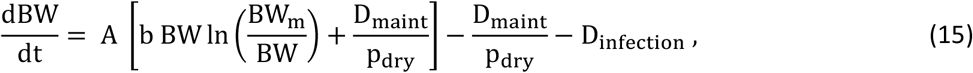

where A, D_maint_, D_infection_ and p_dry_ are specified below in terms of dynamic state variables of the animal. Some previous models of parasite burden include a simplified form of host growth for the purpose of calculating faecal mass and egg output where BW is not related to infection state (e.g. Rose Vineer et al. (2020b); Singleton et al. (2011)), or with a different interaction between growth and parasite burden (Louie et al., 2007). We expect that in a non-infected animal A=1 and D_infection_=0, giving the same gain as for the naïve animal (Eq (11)). In taking resources out of the allocation to growth in Eq. (15), a prioritisation in resource allocation to maintenance and infection functions (Doeschl-Wilson et al., 2008) is implicit.

The DM cost of maintenance can be expressed approximately in terms of metabolic weight, BW^0.75^ (Archer et al., 1997) assuming near thermal neutrality (Supplementary Text S1):

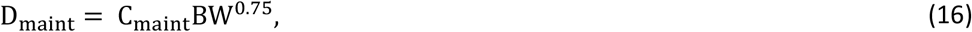

where C_maint_ is a parameter (Table 2). We assume that the costs of infection arise from the increment in the level of immunity (dIm/dt), the maintenance of the level of immunity I_m_ already acquired (Greer and Hamie, 2016), and the repair of damaged tissue associated with the current worm burden:

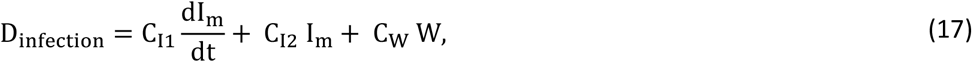

where C_I1_, C_I2_, C_w_ are parameters (Table 2). Values of the parameters that are new, such as C_maint_, C_I1_, and C_I2_ are derived through relationships to other trait parameters reported in the literature (Supplementary Text S2). In this paper we focus on the immunity-related losses and neglect the cost (C_W_=0) of repairing worm-induced damage to the intestine, which is thought to be comparatively smaller (Houdijk et al., 2001).

The DM proportion of gain, p_dry_, is obtained by using an empirically-supported allometric relationship between body water content (Wt) and empty BW (eBW) (Carstens et al., 1991; Filipe et al., 2018), i.e. W_t_=a_w_ eBW^bw^, where a_w_ and b_w_ are allometric parameters, and which, upon differentiation gives

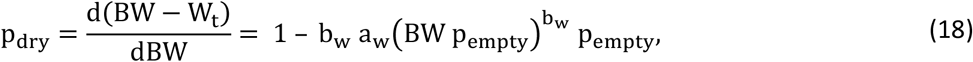

where p_empty_ is the proportion of BW that excludes gastrointestinal tract content (Williams et al., 1992).

The daily feed intake DM (FIDM) is obtained from the FIDDM (Eq. 12 or 13) by accounting for the apparent digestibility of grass DM (DDM) (Colucci et al., 1982):

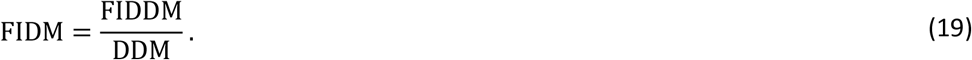

There is evidence DDM is not significantly affected by *O. ostertagi* and other GINs (Fox et al., 1989; Roseby, 1973; Taylor et al., 1989); here, it is assumed to be constant throughout the season (Table 2). Substituting Eq. 12 for FIDDM and Eqs. 14-15 for the remaining terms, Eq. 19 gives

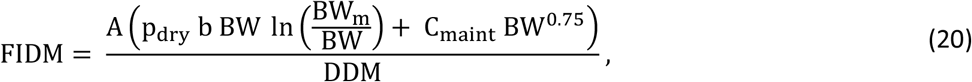

where BW and p_dry_ are given by Eqs. 10 and 18, and the anorexia-related factor A is described below. Note that the term with brackets is the FIDDM of a naïve animal with the same BW. The actual intake was constrained by the capacity of the gastrointestinal tract, which was assumed to be approximately proportional to BW (Text S3).

The daily faecal output that is DM is the total DM intake, FIDM, subtracted of the digestible intake:

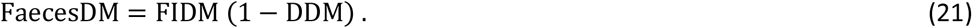

The corresponding daily wet faecal output is:

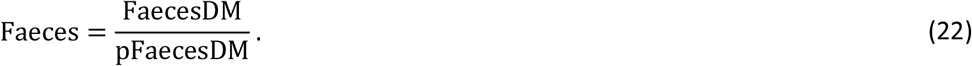

where pFaecesDM is the proportion of faecal output that is DM. For simplicity, pFaecesDM is assumed to be constant at 0.15 throughout the season and across studies (Table 2), although its variability is a known source of uncertainty in FEC observations (Denwood et al., 2012; Le Jambre et al., 2007). The average host FEC (Eq. (6)) is calculated using this dynamically varying prediction of wet faecal output. The parasite-induced reduction in FI is thought to be of the order of 20% to 30% (Bell et al., 1988; Coop et al., 1977; Sandberg et al., 2006); while its causes are not well established (Coop and Kyriazakis, 1999), it is believed to be related to the establishment of new adult worms (Coop et al., 1977), which in the case of *O. ostertagi* occurs in the abomasum (Fox, 1993), where maturation of L4 in the gastric gland provokes inflammation (Charlier et al., 2020a). Therefore, we assumed that A is driven largely by the rate of change in the number of adult worms (dW/dt):

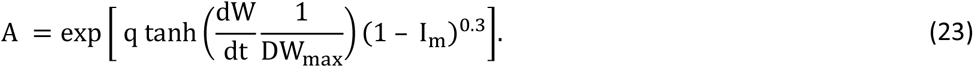

where q = ln(0.8) (Table 2) determines the lowest proportion to which feed intake can be reduced. The function tanh(x), which ranges from −1 to 1, and the scale parameter DW_max_ constrain the effect of dW/dt when its magnitude is of order DW_max_ or greater. When worm load increases (dW/dt>0, tanh(x)>0) then A<1 because q<1; and when the worm load decreases (dW/dt<0, tanh(x)<0) then A>1. Therefore, in the model it is possible to have compensatory growth briefly while the worm number is stabilising, e.g. after the onset of an immune response on worm mortality or after drug treatment. Empirical observations of rebound in BW or FI at a faster pace than may be expected at current BW (Bell et al., 1988; Coop et al., 1977, 1982; Fox et al., 1989; Szyszka et al., 2013) provide suggestive evidence that compensatory growth may occur under these conditions. Such rebounds in appetite and FI, observed rapidly after anthelmintic treatments, are mimicked in the model upon clearing of establishing and adult parasites (Section 2.7), after which parasite-induced anorexia halts, i.e. A=1 as the worm load becomes constant. The immunity-dependent factor in Eq. (23) aims to modulate the magnitude of either effect (A<1 or A>1) when immunity has developed; e.g. when anthelmintic treatment is applied and, following its effects, the worm burden rebounds but immunity has not been lost. The effect of A on FI during compensatory growth was constrained by the capacity of the gastrointestinal tract (Text S3). Other models have made different attempts at incorporating parasite-induced reductions in feed intake based on host variables related to L3 exposure (Berk et al., 2016a; Grenfell, 1988) or adult worm burden W (Louie et al., 2007), while we assumed that change in appetite is driven by change in W.

It has been reported that grazing cattle can have a short-lived drop in BW at the point of turnout caused by a drop in FI and gastrointestinal content (Balch and Line, 1957; Fox et al., 1989) due to adaptation to grazing. We represented this BW drop through a rapid reduction factor in feed intake (Text S4). This correction to intake was applied when modelling empirical studies that exhibited this additional behaviour and when exploring model behaviour.

### 2.4 Sub-model 3: Grass growth

The grass DM available for gazing (G) in a given area of pasture (A_g_), is assumed to be controlled by: the rate of grass growth per ha (r_g_); a carrying capacity per ha (K_g_) that limits grass growth according to characteristics of the grazing system; and the rate of grass intake by the grazing herd at given stocking density (N_h_) and average daily intake per capita FIDM (Table 2). The net rate of grass growth is assumed to have the following growth and consumption terms:

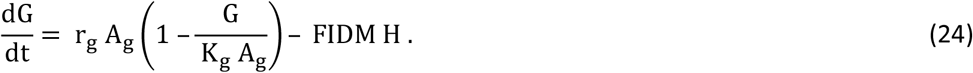

where H=N_h_ A_g_ is the number of hosts in the grazing system. Parameter values are given in Table 3. In this formulation, growth is limited by local resources, i.e. when G/A_g_ approaches K_g_ growth stops and any further grazing will lead to decrease in sward availability, as is observed (Dimander et al., 2003; Larsson et al., 2006). The assumed value of G at turnout (Table 3) is such that the grazing system has not yet reached its limit capacity; hence, some increase in G (dG/dt>0) is possible upon moderate consumption. Equation (24) assumes that the grass plants are only increasing in size and not propagating in number; the opposite assumption can be made through logistic growth, where the growth term in Eq. (24) would be multiplied by G (Grenfell, 1988; Louie et al., 2007). In using empirical measurements of r_g_ (Table 3) we have assumed that these were obtained without, or were discounted for the latter density effects, which is an approximation. Other authors have chosen not to include such limiting effects in grass growth (Berk et al., 2016b). The rate of grass growth and the carrying capacity are currently assumed to be constant throughout the season. In addition, in the current non-spatial formulation, all variables are assumed to be uniform across the grazing area: the herd grazes an evenly-distributed herbage and each animal grazes identically. The grass availability G/Ag is used to convert the density of L3 per ha into the concentration of L3 per kg DM (Table 1), used to calculate the ingestion of L3 per animal.

### 2.5 Sub-model 4: Free-living stages

A model of the dynamics of the parasite’s FL stages has been fully developed previously (Rose et al., 2015). Building on past work (Grenfell et al., 1987a; Smith, 1994), the GLOWORM-FL model incorporated the migration of infective L3 larvae between soil and herbage and the influence of weather variables on this movement. The model also contained fresh estimation of the influence of weather on the remaining parameters controlling the FL stages. In the current paper, we have added to this sub-model two dynamic flows linking the nematode FL and parasitic stages: the deposition of eggs in faeces and the ingestion of L3 by every host in the grazing herd. We briefly describe the model’s variables and parameters (Table 4) and the additions to the model. Details, including parameter values for *O. ostertagi* are given in Text S4.

The dynamics of the FL living stages, including deposited eggs (E_p_), stages developed within faecal pats (L_12_, L_3f_), and L3 on pasture (L_3p_), on herbage L_3h_, and on soil (L3s), are defined as densities per ha and given by the rate equations (Rose et al., 2015):

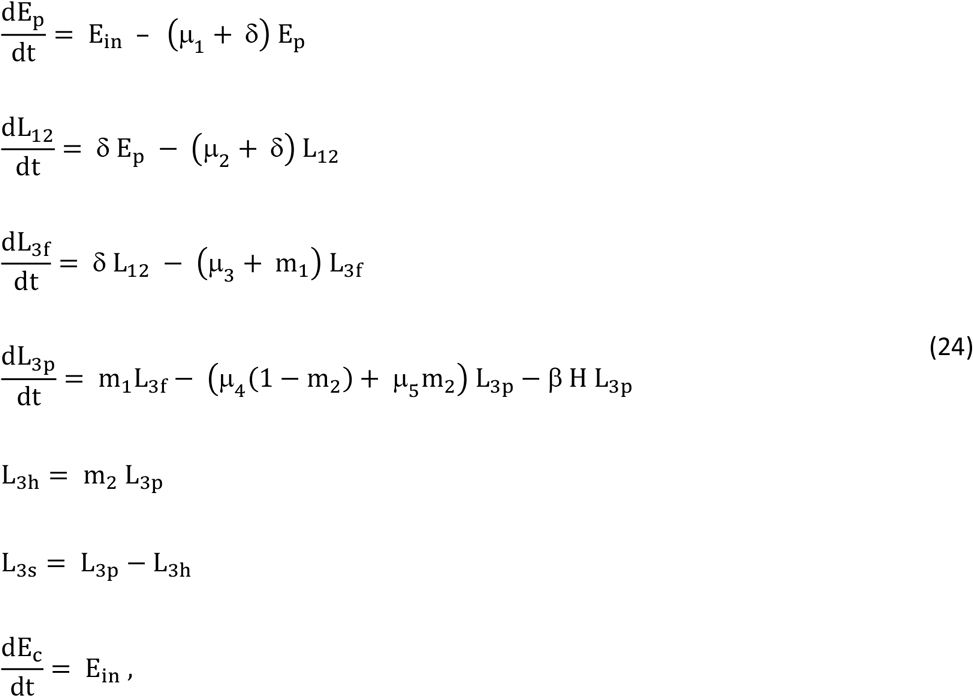

All variables and parameters are described in Table 4. The assumed initial values of the variables at turnout are given in Supplementary Table S2. The last equation in Eq. (24), for the cumulative number of eggs deposited (Ec), was introduced by us for later use. The remaining equations are as in Rose et al. (2015), but with three exceptions. First, we have replaced the rate δ for 2δ in Rose et al. (2015). Second, the rate of egg deposition on pasture by all hosts per day per ha was 100 and is now replaced with:

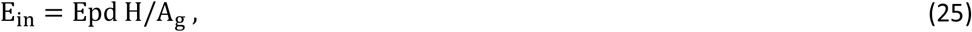

where Epd, H and A_g_ are as in Tables 1 and 3. Third, there is an additional term in the rate of change of L_3p_ representing the daily ingestion of L3 by every grazing animal per ha. This term required defining a new time-varying parameter. The average daily probability of ingestion per L3 per host (β(t)), known as rate of parasite transmission or instantaneous rate of infection, is the ratio of the L3 ingested per grazing host per day per ha (FIDM m_2_ L_3p_/G) to the L3 available on pasture per ha (L_3p_):

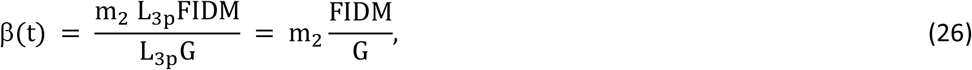

where FIDM (Table 2, Eq. 20) is determined by the host’s BW, parasite burden and level of immunity; G (Table 3, Eq. 23) is the current grass biomass available for grazing; and m2 (Table 4, Eq. 24) is the current weather-dependent availability of L3 on herbage. The ingestion term, β H L_3p_, in Eq. 24 is analogous to the transmission term in other models (Grenfell, 1988; Grenfell et al., 1987a; Kao et al., 2000; Louie et al., 2005; Roberts and Grenfell, 1991; Singleton et al., 2011; Smith and Grenfell, 1985); the difference being in how β is defined, which we do in terms of a host state variable, grass mass, and environmental drivers. Other work defined β conceptually similarly in terms of constant quantities (Singleton et al., 2011), through the ratio FIDM/G with FIDM determined by grass mass and reduced by larval exposure (Grenfell, 1988), or as a function of host age (Louie et al., 2005). Predictions of β are provided later. Note that Eq. 26 feeds into Eq. 1, which closes the loop of interdependency of model variables.

### 2.6 Reproduction number

To characterise the increase or decrease of the parasite population, and thus whether it is controlled, we quantify the average extent to which each individual parasite replaces itself during its lifetime. For macroparasites, the basic reproduction number (R_0_) is defined as “the average number of (female) offspring per adult (female) worm that survive to reproduction in the absence of density-dependent constraints” (Anderson and May, 1992; Tompkins et al., 2001). Heuristic (Anderson and May, 1992) and formal (Heesterbeek and Roberts, 1995) calculations of R_0_ have been provided for simple models of parasites with direct cycles; they quantify the parasite’s maximum replacement rate when its inherent reproduction and survival traits are expressed to the full extent, typically early in an outbreak. Here, we focus on the overall dynamics of the parasite population towards stability by considering the effective reproduction number (R_e_), which includes the regulatory effects imposed by the host and the environment on each parasite stage (Churcher et al., 2006). Given that it is not straightforward to calculate R_e_ amid the complexities of the current model (Filipe et al., 2005), R_e_ can be expressed via the following time-varying factors:

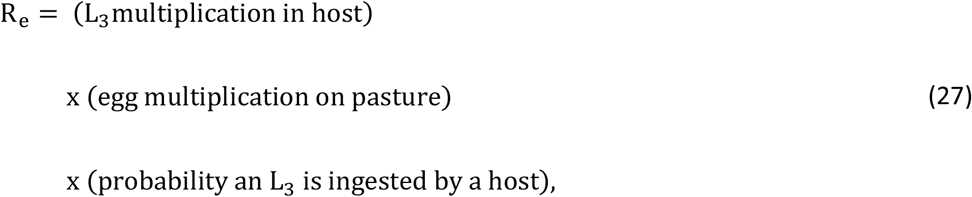

Each of these factors can be expressed approximately and respectively as:

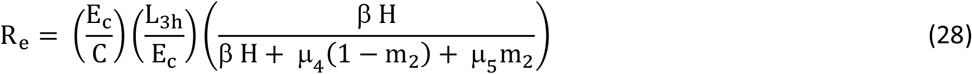

In the first factor, E and C are the cumulative numbers of eggs produced and L3 ingested by a host; in the second factor, L_3h_ and E_c_ are the cumulative numbers of L3 on herbage and eggs laid on pasture; and the third term is the average proportion of L3 on pasture ingested by hosts, given by the ratio of the rate of L3 ingestion per day per ha (β H, Eq. 26) to the rate of L3 departure from pasture through ingestion or mortality per day per ha (β H + μ_4_ (1- m_2_) + μ_5_ m_2_). This calculation is heuristic and approximate in its use of ratios of cumulative numbers of outgoing to incoming parasites per stage. The cumulative aspect tackles the fact that, under time varying conditions, changes in incoming and outgoing parasite stages are not synchronous; as it would be difficult to incorporate time lags explicitly, the calculation is approximated through the use of time averages. In a parasite population that stabilises, we would expect R_e_ to become close to 1, indicating no increase or decrease. However, full stabilisation through the regulatory factors that reduce R_e_ may take time to unfold on the scale of a single season. In addition, there is variation in environmental conditions due to seasonal climate, weather and management actions likely to cause fluctuations before and after stabilisation. Nonetheless, a R_e_ that declines over time to magnitudes around 1 would provide a health check on the mutual consistency of the parameters of the host and free-living sub-models.

### 2.7 Model behaviour

#### 2.7.1 Numerical implementation

The model was solved numerically using Euler’s method with a time step of 0.1 day. This step is small enough at the scale of all processes represented in the model and thus is likely to lead to solutions with satisfactory numerical accuracy. Using a step smaller than 0.1 led to no observable difference in the model output. In addition, this accuracy was assessed on simpler models with known analytical solution, giving an acceptable relative error of 0.37% with step 0.1 d, 3.6% with step 1d, and 28% with step 10d. The model was coded in the R language and the results were generated using the free software R, version 4.1.1 (R Core Team, 2021. R: A language and environment for statistical computing. R Foundation for Statistical Computing, Vienna, Austria). A code of the model is available (Filipe, 2022).

#### 2.7.2 Baseline system

Predictions of the model, as defined in Sections 2.2-2.5 using parameter values from literature, were validated using the approach in Section 2.8. In addition, we explored how the model captures the effects of key factors on the epidemiology and control of *O. ostertagi*. For this purpose, we used baseline conditions defined by a representative location of temperate weather in Northern Europe and a typical year among its records of daily weather. We chose as location Large Park Hillsborough, BT26 6DR in Northern Ireland (coordinates 54°27’06.6’’N, 6°04’30.7’’W). Weather data (daily mean temperature and total precipitation) for this location were collected from the E-OBS gridded dataset (Cornes et al., 2018). We chose 2014 as a typical year among the last 10 years of weather data (2011-2020) as the daily pattern and annual average of the temperature in 2014 were closest to those of the daily records averaged over 10 years. As a representative grazing period we used 01/05 to 25/09 (21 weeks) in 2014. Weather data were used raw, without smoothing. The weather variables are plotted in Supplementary Fig. S1.

The baseline parasitology at turnout had an average concentration of *O. ostertagi* on herbage of 200 L3/kgDM (Berk et al., 2016b), and assumed that no other FL stage overwintered (Supplementary Table S2). We note that in the model this is the actual level of L3 on herbage, while in a real system an observation of L3 on herbage is likely to be an underestimation of its actual level; e.g. an assumed level of 200 could correspond to an observed level of 100 or less. The baseline FGS calves were assumed to be naïve (parasite free and with no acquired immunity), to have a body weight of 200 kg at turnout and growth parameters as in Table 2, which led to a BW trajectory in the range 200-400 kg over the 21-week grazing period. Daily FI was assumed to drop at turnout (Section 2.3 and Supplementary Text S4) as observed in some of the empirical studies used for validation and often observed more generally (Balch and Line, 1957). The cattle herd was assumed to have a stocking density of 5 animals/ha (Table 3).

#### 2.7.3 Behaviour explored

Assuming the model structure and the parameters values described in Sections 2.2-2.5, the behaviour of the baseline system was explored in a range of scenarios where one model parameter was varied at a time:

1. Effect of the initial level of herbage contamination, i.e. concentration of L3 on herbage at turnout: L_3c_ = 100, 200, 500 L3/kgDM (Michel et al., 1970).
2. Effect of the herd stocking density: N_h_=1, 5, 7 animals/ha, where 1-5 is the range found in the empirical studies used for model validation.
3. Effect of one anthelmintic treatment differing in the timing of application: T1 = 0, 4, 8 weeks after turnout.
4. Effect of two anthelmintic treatments with the first treatment applied at turnout (T1=0) but differing in the time of application of the second treatment: T2 = 3, 5, 7 weeks from turnout.

A simplified drug treatment was modelled, with 100% efficacy in clearing establishing and adult parasite stages during 21d and with no effect afterwards. The variation of some grass-growth parameters (Table 3) was also explored but found to have limited influence on model behaviour under the current values of the other parameters. These explorations were not included in the results but confirmed that our choice of values for these parameters was not determining.

### 2.8 Model validation

#### 2.8.1 Datasets

In order to test the predictions of the model defined in Sections 2.2-2.5, the literature was searched for empirical studies satisfying the following criteria:

1. The study was on FGS beef or dairy cattle (aimed at testing the model on predominantly naïve animals).
2. Longitudinal data were provided on BW and FEC (the most common field observations) from turnout until the end of the experiment; where other parasitological variables predicted by the model were reported these were also compared with the model predictions.
3. Artificial dosing with L3 larvae was not used during the course of the study or was used only at the point of turnout (aimed at allowing infections to occur naturally after completion of the parasite’s full cycle through the environment).
4. The experiment contained a group of untreated animals (aimed at testing the model in the absence of anthelmintic treatment, and to ensure that treatment applications in the model overlay a plausible host response).
5. The experiment contained an additional group of animals treated prior to and after the point of turnout (aimed at using the weight data of this group as proxy data for the weight of non-infected animals); for this purpose of characterising growth the animals should be genetically similar, i.e. from a single breed or cross of breeds.
6. The study location and calendar dates were provided so that weather data for the duration of the experiment could be obtained.
7. The study location was in Northern Europe.

We identified six studies based on these criteria (Table 5). Studies with natural infections only: 1) Larsson et al. (2007); Larsson et al. (2006), and 2) O’Shaughnessy et al. (2015). One study where the animals were inoculated with lower doses of L3 at turnout: Höglund et al. (2018), comprising 1) dairy cattle and 2) crossbred cattle from dairy and beef breeds. Studies where the animals were inoculated with higher doses of L3 at turnout: 1) Dimander et al. (2003) and 2) Höglund et al. (2013). All studies took place either in Ireland or in Sweden. Data were available from tables, text or figures in each article.

**Table 5.** Six empirical studies used for model validation. Summary of the information provided.

| Study | Location | Year | Duration (d) | Av. BW at turnout (kg) | Av. L3 on pasture at turnout (1/kgDM) | Dose at turnout (L3) | Stock. dens. (1/ha) | Additional measures | Co-infection *O. ostertagi* (%) vs *C. oncophora*, serial observations |
| --- | --- | --- | --- | --- | --- | --- | --- | --- | --- |
| O'Shaughnessy et al. 2015 | Ireland | 2012 | 124 | 165 | 200<sup>1</sup> | - | 1.4 | L3c (SGS) | L3: 73%, 23%, 50% (SGS) |
| Larsson et al. 2006; 2007 | Sweden | 2002 | 148 | 189 | 250<sup>2</sup> | - | 5 | L3c (Y1, Y2), W (tracers 3w prior housing) | L3:<50% all season; W: 30% |
| Höglund et al. 2018, dairy | Sweden | 2016 | 142 | 306 | <sup>3</sup> | 5000<sup>4</sup> | 2.25 | - | L3: 47%, 80% (PCR) |
| Höglund et al. 2018, cross | Sweden | 2016 | 142 | 332 | <sup>3</sup> | 5000<sup>4</sup> | 2.25 | - | L3: 17%, 47% (PCR) |
| Höglund et al. 2013 | Sweden | 2008 | 154 | 238 | <sup>3</sup> | 40000<sup>4</sup> | 2.4 | W (tracers Y2, Y3) | W: 24%, 27% |
| Dimander et al. 2003 | Sweden | 1999 | 150 | 200 | 300 | 10000<sup>4</sup> | 5 | L3c (Y1, Y2); W (Y4 tracers) | L3: 20%, 75% |

<sup>1</sup> Based on SGS.
<sup>2</sup> Average of first two observations.
<sup>3</sup> No data provided (c.f. note in Section 2.7.2).
<sup>4</sup> Doses comprised equal proportions of *O. ostertagi* and *C. oncophora*. FGS, SGS: first and second grazing season. Y1, Y2: Year 1, Year 2.

All studies reported mixed infections comprising predominantly *O. ostertagi* and *C. oncophora* (Table 5). The model, which was designed for a single-species *O. ostertagi* infection, was compared with these data as we lacked single species data.

#### 2.8.2 Validation approach

For each empirical study, we compared the model predictions with the longitudinal observations of BW (or gain plus average start weight, as provided), and FEC. These data were reported as averages over the animals in each treatment group and at each time point from turnout to the end of the experiment. Where available, we also compared with model predictions any L3 observations throughout the experiment, and worm counts in tracer animals from within the untreated group or that grazed the same paddocks in subsequent seasons. There were no data on feed intake. Weather data (daily mean temperature and daily total precipitation) for the spatial coordinates and calendar dates reported in each study were collected from the same source as the baseline weather and used raw, without smoothing. The weather variables are plotted in Supplementary Fig. S1.

The local daily weather, initial L3 contamination of herbage, any inoculation dose at turnout, cattle stocking density, and cattle breed growth parameters were the only quantities adjusted to describe the conditions of each empirical study. While many other parameters could have differed among studies, all other model parameters were assumed to be the same across all studies, i.e. there was no model fitting to data. In one study (O’Shaughnessy et al., 2015), the FEC at turnout was positive, hence it was necessary to assume initial non-zero values for the number of adult worms and immunity level.

Where no measures of L3 on herbage at turnout were available (Table 1), an initial herbage contamination was assumed based on studies in comparable regions (Michel et al., 1970) following a similar reasoning as for the baseline system. Using the model’s predicted mean trajectory of L3 concentration on herbage, we drew a sample from a negative binomial distribution (Smith and Guerrero, 1993) with this mean and an aggregation parameter k=1.4 (Verschave et al., 2015); the samples’ lower quartile was contrasted with the data in an attempt to account for low efficacy in the field recovery of L3 (Kloosterman, 1971; Paras et al., 2018). As the breed of the animals differed across studies and their growth parameters were unknown, the average BW of the group of treated animals was used as proxy for the BW of a naïve animal of the same breed, which unlike the infected animals is not affected by anorexia and infection costs; these BW data were used to estimate the performance parameters of the Gompertz BW gain of the infected animals (Eq. 11). This estimation was derived using the nonlinear model regression function nls of the software R, version 4.1.1 (R Core Team, 2021. R: A language and environment for statistical computing. R Foundation for Statistical Computing, Vienna, Austria). Standard errors for BW and FEC mean observations were provided in a minority of studies and included in the plots where the model output and empirical data were compared.

To be able to tackle data from two-species co-infections (Section 2.8.1 and Table 5) and which do not specify parasite numbers by species (except for occasional relative proportions in some studies), we input the initial total concentration of L3 on herbage into the model and predicted the numbers of parasitic and FL nematode stages as totals for both species, which we then compared with the data.

The working hypotheses are that: 1) the parameters and processes in the model are adequate for describing each species and thus the total infection, and 2) there are no interactions between species. In this way, the model is regarded as representing a typical parasite mixture with variable relative proportions of *C. oncophora* and *O. ostertagi.* Alternatively, if we knew the proportions of observed FEC and initial L3 corresponding to *O. ostertagi*, we could have modelled a single *O. ostertagi* infection (Smith and Guerrero, 1993), which would nevertheless still assume no interactions between species in the real system. Unfortunately, these proportions vary throughout the season (Högberg et al., 2021) and are largely unknown as indicated by the rare measurements in the current studies (Table 5); this was the case whether the animals were inoculated with known species mixtures at turnout or subject solely to natural infection. For these reasons, we considered inevitable to take the above approach, which is simple and easily interpretable.

#### 2.8.3 Statistical approach

A statistical comparison between the BW and FEC predicted by the epidemiological model and the empirical data was made using a standard validation approach (Mayer and Butler, 1993). In this approach, the observed data are linearly regressed on the model output, i.e. the first is treated as a response and the latter as a predictor; the intercept of the relationship is fixed at zero. The outcomes of this regression are an estimated slope, a p-value on an F statistic assessing the fitted line against a constant response, a 95% confidence interval (CI) on the estimated slope, and a coefficient of determination adjusted for the number of model parameters (R^2^adj). If the p-value is significant, we can reject the null hypothesis that there is no relationship between the observations and the model (i.e. that there is no change in the observations the predictions change); we used the conventional 5% significance level, but expect that only much smaller p-values would comfortably reject the null hypothesis. If the 95% CI of the estimated slope includes the value 1, in addition to excluding the value 0, then there is statistical support for the epidemiological model as this indicates the overall deviation between model and data is within the variation expected to occur within the data. An R^2^_adj_ close to 1 supports the assumed linear relationship between the epidemiological model and the data. We report the p-value, CI on the slope, and R^2^_adj_. The statistical analyses were done using the linear model regression function lm of the software R, version 4.1.1 (R Core Team, 2021. R: A language and environment for statistical computing. R Foundation for Statistical Computing, Vienna, Austria).

## 3. Results

### 3.1 Model behaviour

The response of the model to differing conditions was explored using the model structure and parameters of Sections 2.2-2.5. Further results, on the validation of the model, are presented in Section 3.2.

#### 3.1.1 Effect of the initial level of herbage contamination

Varying the herbage contamination at turnout (L3c = 100, 200, 500 L3/kgDM) caused a considerable peak shift (earlier peak) in calf FEC and in adult worm burden (Fig. 2). A higher starting pasture contamination caused an earlier, higher peak and an earlier decline in FEC. For the adult worm burden, however, a higher initial contamination caused an earlier but lower peak and a later decline; this difference can be due to the fact FEC is affected by additional and earlier effects, i.e. reduction in fecundity due to acquired immunity and density dependence. The effects on BW were appreciable, with greater differences in cumulative gain between 10-15 weeks, but were followed by some recovery in lost gain and more modest BW differences by the end of season. The differences in contamination at turnout were reflected in the temporal trajectories of L3 contamination, but were much reduced by the end of the season.

**Fig. 2.**
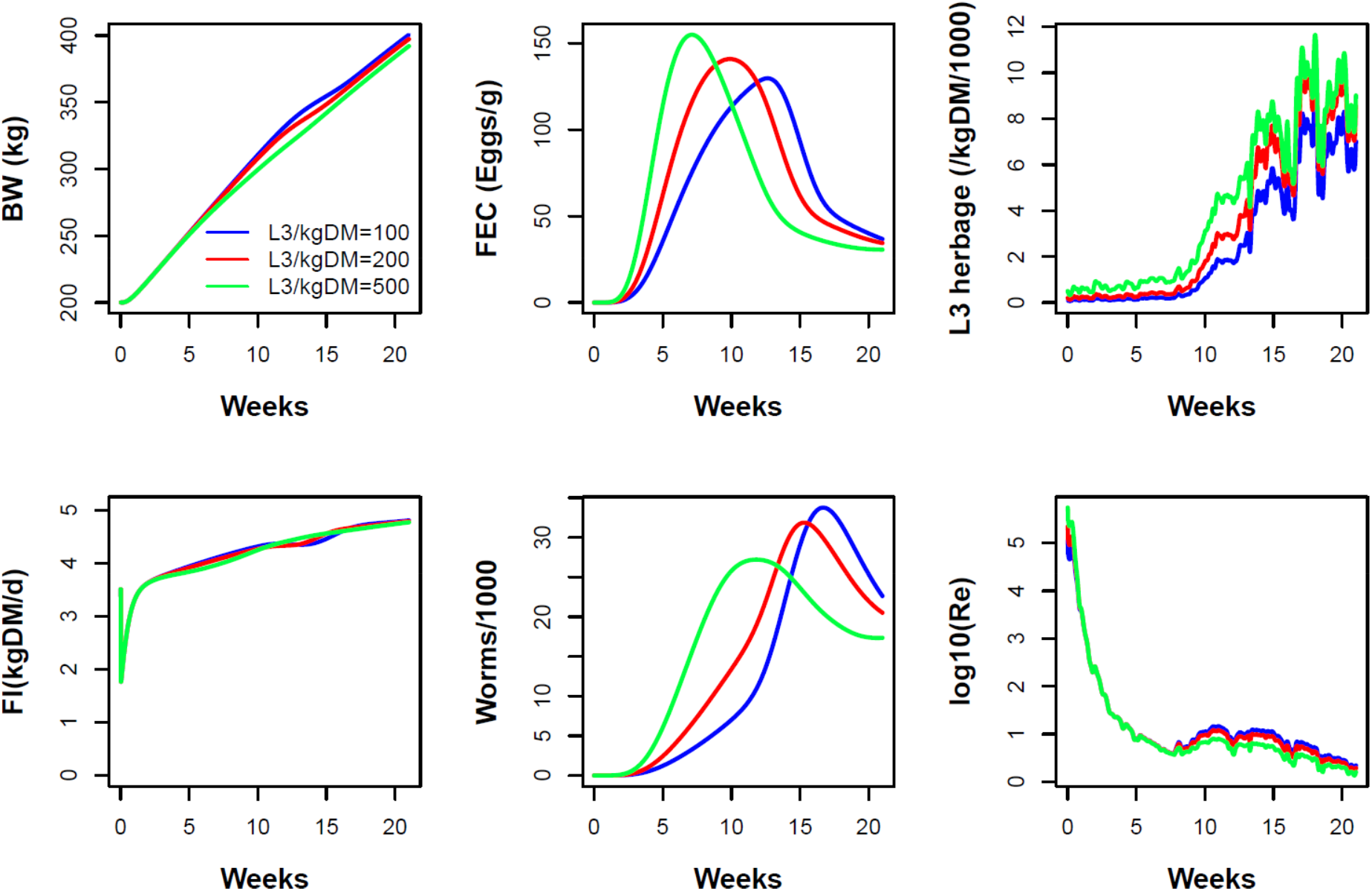
Model behaviour: effect of the initial level of herbage contamination. Progression of *O. ostertagi* infection during the grazing season of the baseline herd and grazing system under differing initial concentrations of L3 on herbage, 100, 200, 500 larvae/kg dry matter (DM). Traits shown: body weight (BW), faecal egg count (FEC) in wet faeces, density of L3 on dry herbage, daily feed intake (FI), number of adult worms, and logarithm of the effective reproduction number R_e_.

#### 3.1.2 Effect of the herd stocking density

Higher stocking densities (N_h_=1, 5, 7 ha^-1^) enhanced transmission and amplified infection pressure, leading to earlier, higher peaks in worm burden and FEC, which then declined towards the end of the grazing season (Fig. 3). At lower stocking densities, egg shedding from animals persisted longer due to lower levels of immunity. However as stocking density increased, pasture contamination increased markedly and persistently, as a result of a larger number of hosts shedding eggs and infective larvae developing from earlier, higher shedding. In addition, the greater parasite challenge at high stocking densities resulted in slower weight gain that produced persistent differences in body weight through to the end of the grazing period (Fig. 3). Across weather in different years, these patterns were robust but the strength of peak shifts in FEC and differences in BW by the end of season vary across years (not shown); e.g. these effects were stronger in 2011 than in the current example of 2014. Compared with increases in the initial pasture larval contamination, increases in stocking density drove later but more persistent differences in FEC, herbage larval level, and weight gain; and, for worm burden and FEC, differences mostly in magnitude rather than timing. These differences result from the time lag between host infection, through egg shedding to larval maturation, and from the greater number of hosts carrying and transmitting parasites. Note that the red curves are identical between Figs. 2 and 3. Increased L3 abundance late in the season at higher stocking densities (Fig. 3) could affect starting L3 levels in the next season (Fig. 2).

**Fig. 3.**
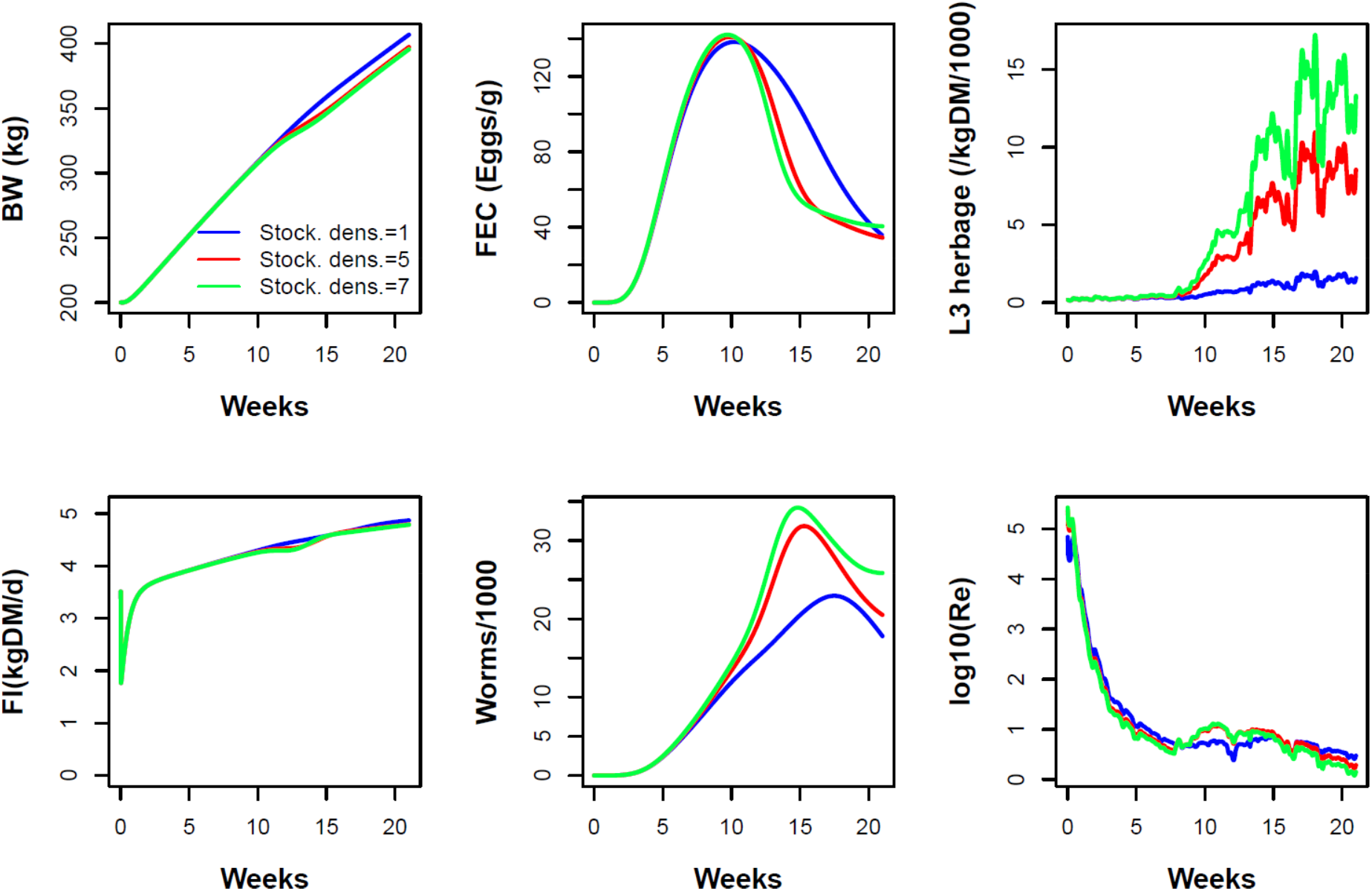
Model behaviour: effect of the herd stocking density. Progression of *O. ostertagi* infection during the grazing season of the baseline herd and grazing system under differing herd stocking densities, N_h_ = 1, 5, and 7 animals/ha. Traits shown: body weight (BW), faecal egg count (FEC) in wet faeces, density of L3 on dry herbage, daily feed intake (FI), number of adult worms, and logarithm of the effective reproduction number R_e_.

#### 3.1.3 Effect of one anthelmintic treatment differing in the timing of application

A key management parameter when a single round of drug treatment is applied after turnout, is the timing of application during the grazing season. The model predicted that an intermediate time after turnout (among 0, 4 or 8 weeks post turnout) is optimal (Fig. 4) in the following sense: it led to greater cumulative BW gain and to lower cumulative parasite burden in the host, and thus to potentially lower risk of production loss and clinical disease, while still leading to end-of-season pasture contamination and parasite burden comparable to those of the late treatment. Early treatment (at turnout) delayed infection but also immunity, leading to higher overall worm burdens and late-season L3 levels than for the later treatments, although still lower than in the absence of any treatment (Fig. 3, red line).

**Fig. 4.**
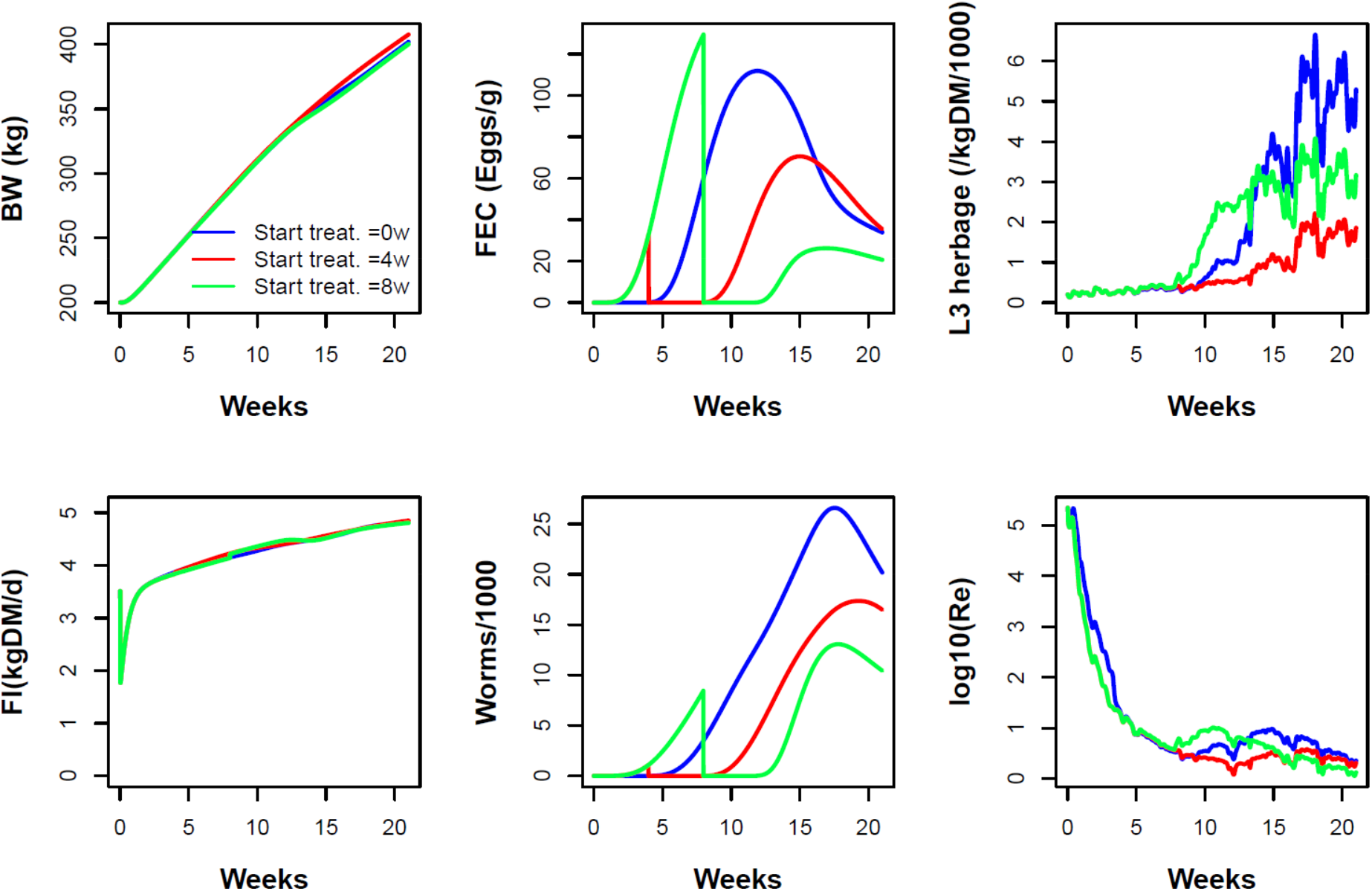
Model behaviour with one anthelmintic treatment: effect of the timing of application. Progression of *O. ostertagia* infection during the grazing season of the baseline herd and grazing system with one round of anthelmintic treatment with differing times of application, 0, 4 and 8 weeks after turnout (see Section 2.7.3 for details on drug treatment). Traits shown: body weight (BW), faecal egg count (FEC) in wet faeces, density of L3 on dry herbage, daily feed intake (FI), number of adult worms, and logarithm of the effective reproduction number R_e_.

#### 3.1.4 Effect of two anthelmintic treatments differing in the timing of the second treatment

When two anthelmintic drug treatments are applied, it is natural to apply the first at turnout and to examine the best timing for application of the second treatment. The model predicted that when varying the latter (among 3, 5, 7 weeks post turnout), there was not much difference in the final BW (Fig. 5). In all cases, the lowered infection pressure led to lower levels of immunity, which allowed adult worm populations to continue increasing throughout the grazing period. However, there was considerable difference in the level of pasture contamination potentially carried over to the next season and in the cumulative parasite burden (Fig. 5), with the latest application being the best in this respect, but with the intermediate timing being next best.

**Fig. 5.**
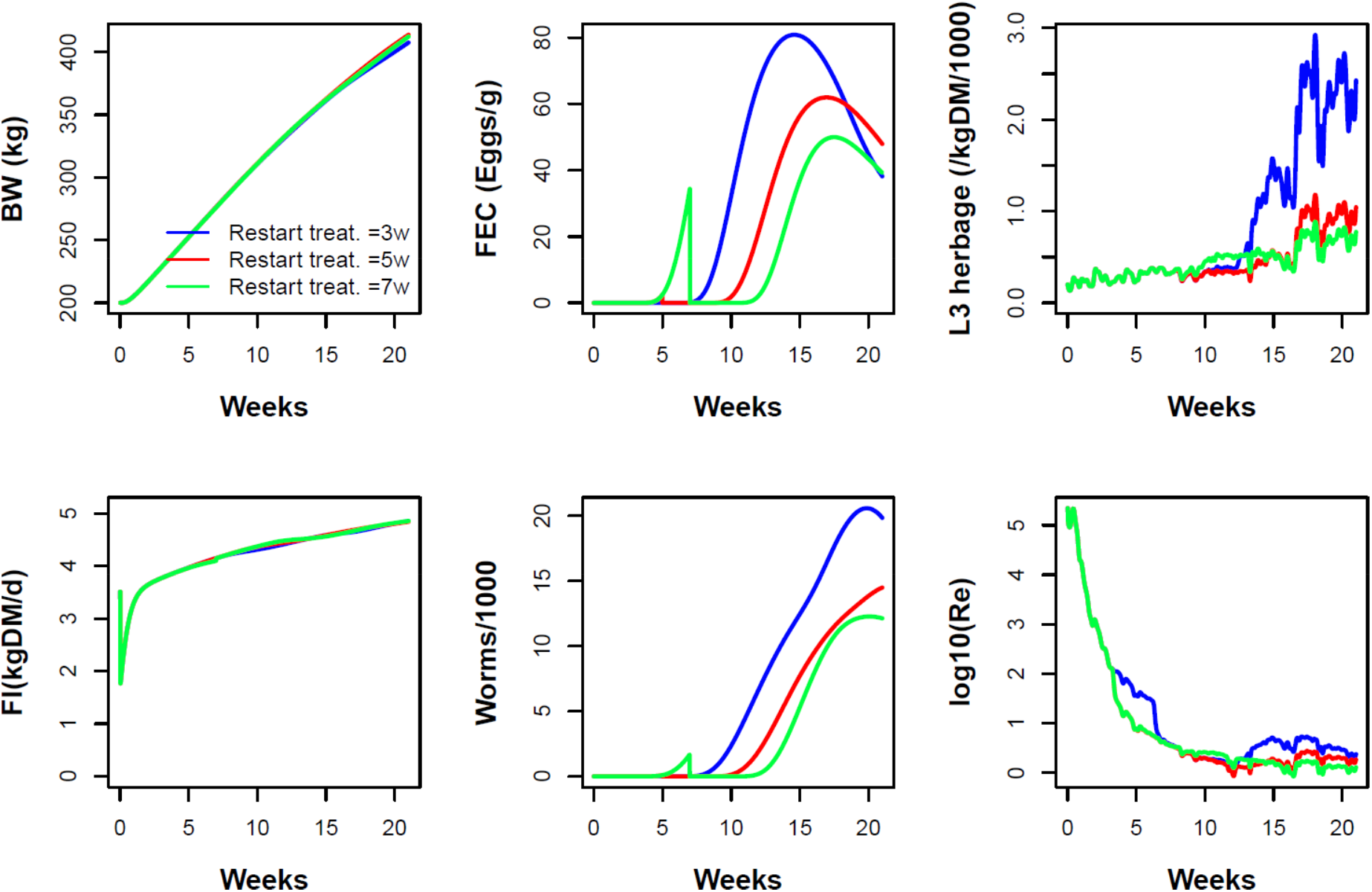
Model behaviour with two anthelmintic treatments: effect of the timing of the second treatment. Progression of *O. ostertagi* infection during the grazing season of the baseline herd and grazing system with a first round of anthelmintic treatment at turnout but differing in the time of application of the second round, at 3, 5, and 7 weeks after turnout. Drug efficacy is maintained for 3 weeks (see text for details on drug treatment). Traits shown: body weight (BW), faecal egg count (FEC) in wet faeces, density of L3 on dry herbage, daily feed intake (FI), number of adult worms, and logarithm of the effective reproduction number R_e_.

#### 3.1.5 Effective reproduction number

In the four model behaviour examples, the predicted effective reproduction number R_e_ exhibited a clear pattern (shown on a logarithmic scale, Fig. 2-5). In the initial phase R_e_ had very large values likely influenced by the assumed initial numbers of parasite stages and by specifics of the calculation of R_e_ (Eq. (28)). Hence, during this phase, we seek for meaning in pattern rather than in values. Subsequently, there was a sharp decline in R_e_ followed by oscillations within the range 10-1, i.e. above but not far from the value 1. This pattern confirms our expectations about the model output (Section 2.6): it indicates convergence to a state of quasi stability in the parasite population superimposed by short-term fluctuations, likely due to variation in host response and weather (the same in the four examples). In these examples, the curve with lowest final value of R_e_ does not always correspond to the case with the lowest final value of L3; we expect this to be because R_e_ is also affected strongly by parasite burden and the current level of feed intake.

### 3.2 Model validation

Predictions of the model defined in Sections 2.2-2.5 were tested against data from empirical studies. The studies were organised in pairs that had, respectively, natural infection (Fig. 6), a low artificial parasite dose followed by natural infection (Fig. 7), and a higher artificial parasite dose followed by natural infection (Fig. 8). These figures show the predicted BW, daily FI, FEC, and number of adult worms per FGS calf, and the predicted L3 herbage contamination and R_e_. The figures also show the observations on BW and FEC and, where data were available, on other traits.

**Fig. 6.**
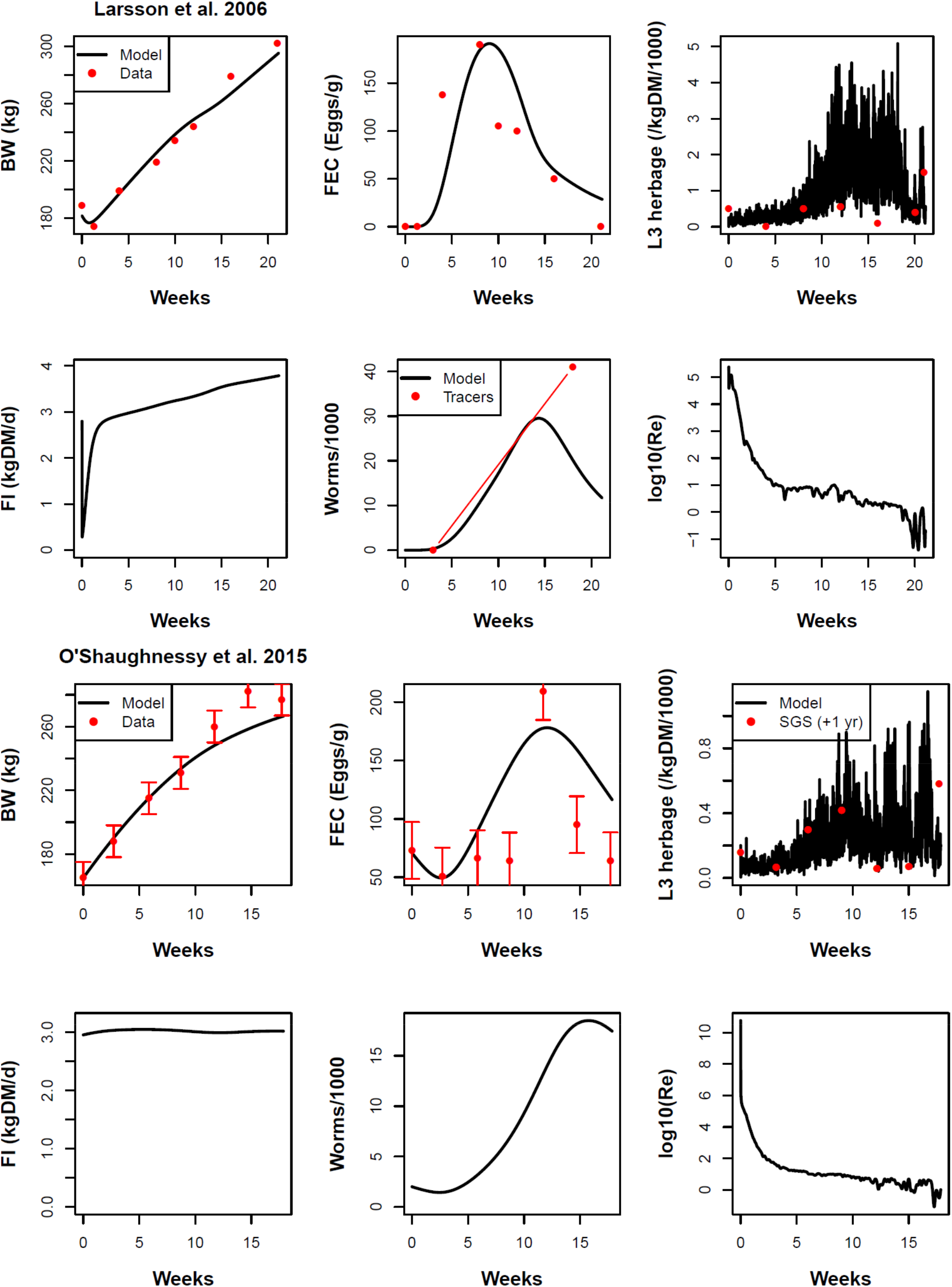
Model comparison with studies involving natural infection of cattle: Larsson et al. (2006) (rows 1 and 2) and O’Shaughnessy et al. (2015) (rows 3 and 4). Traits shown: body weight (BW), faecal egg count (FEC) in wet faeces, density of L3 on dry herbage, daily feed intake (FI), number of adult worms, and logarithm of the effective reproduction number R_e_. Animals were infected with a mixture of *O. ostertagi* and *C. oncophora.* The results of the statistical tests are given in Table 6.

**Fig. 7.**
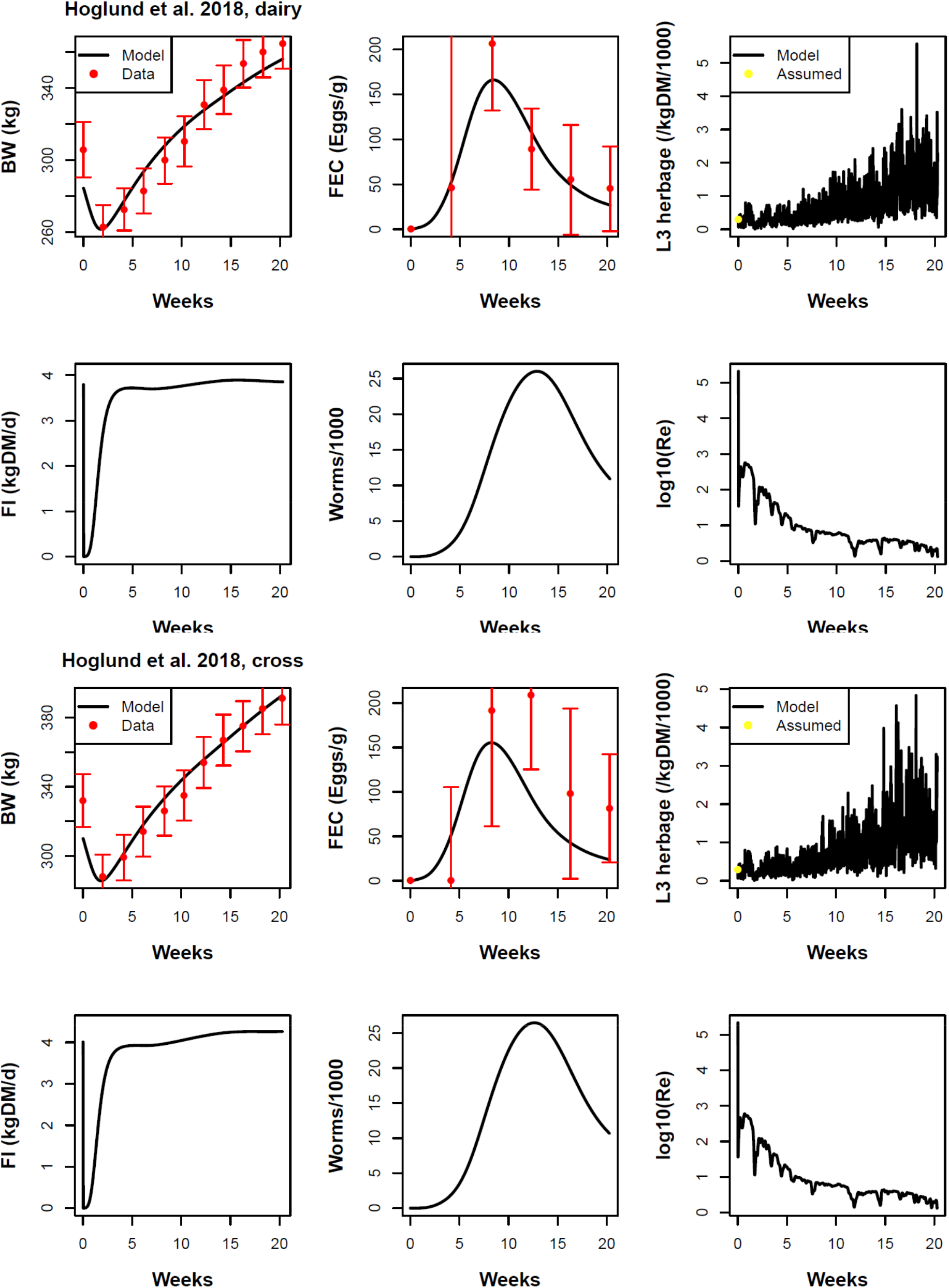
Model comparison with studies providing a low parasite dose pre-turnout followed by natural infection: Höglund et al. (2018) dairy breed (rows 1 and 2) and cross breed (rows 3 and 4). Traits shown: body weight (BW), faecal egg count (FEC) in wet faeces, density of L3 on dry herbage, daily feed intake (FI), number of adult worms, and logarithm of the effective reproduction number R_e_. The parasite dose at turnout was a 5000 even mixture of *O. ostertagi* and *C. oncophora.* The results of the statistical tests are given in Table 6.

**Fig. 8.**
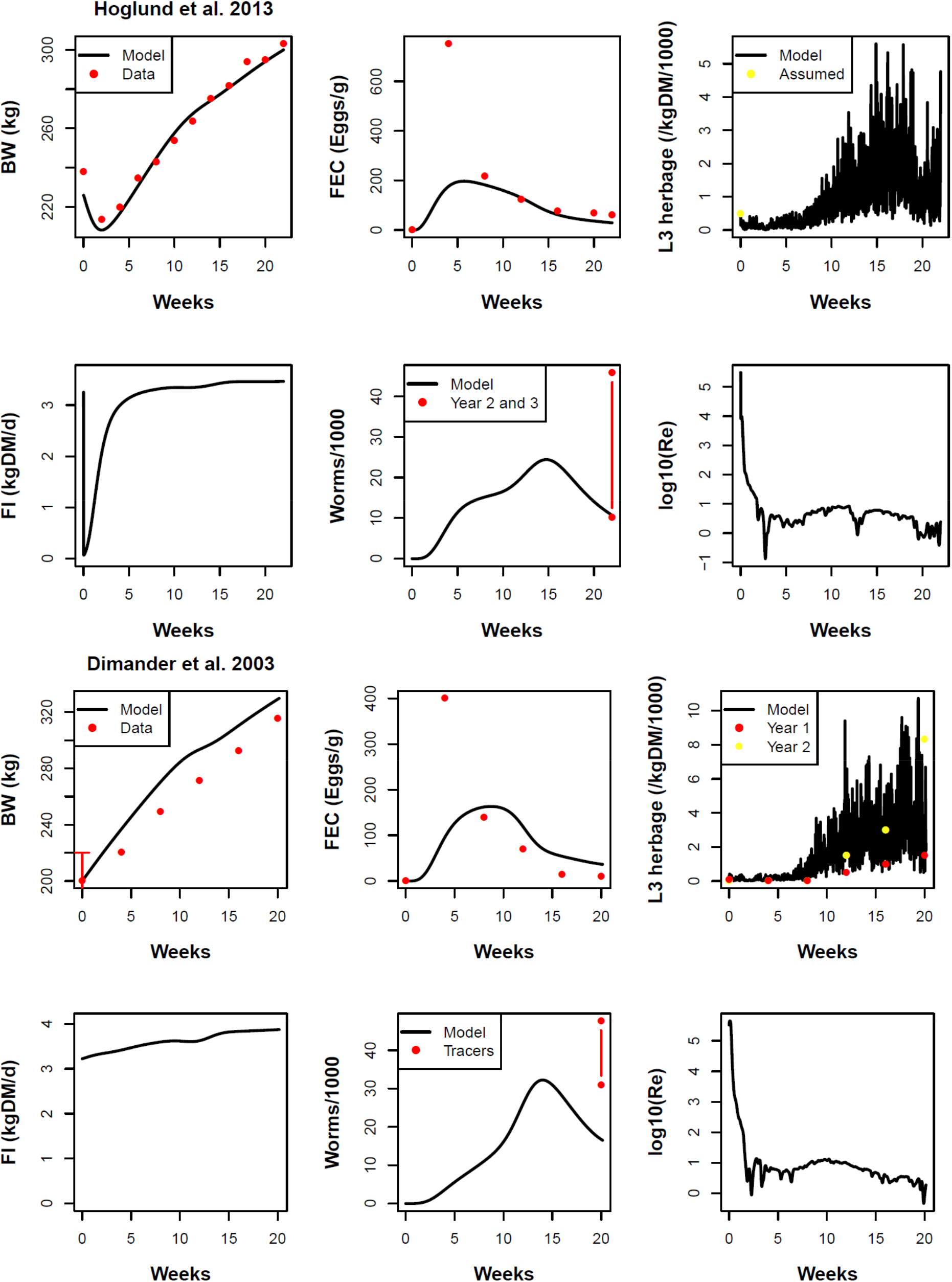
Model comparison with studies providing a higher parasite dose pre-turnout followed by natural infection: Höglund et al. (2013) (rows 1 and 2) and Dimander et al. 2003 (rows 3 and 4). Traits shown: body weight (BW), faecal egg count (FEC) in wet faeces, density of L3 on dry herbage, daily feed intake (FI), number of adult worms, and logarithm of the effective reproduction number R_e_. The parasite dose at turnout was a 40000 (top) and 10000 (bottom) even mixture of *O. ostertagi* and *C. oncophora.* Results of the statistical tests are given in Table 6.

#### 3.2.1 BW and FEC

Visual comparison of the predictions with the empirical data indicates the model was generally in reasonable agreement with the BW and FEC data across the studies (Fig. 6-8). There is formal statistical support for this agreement in almost all cases (Table 6): the 95% CI of the slope of the relationship between observed data and prediction includes the value 1 and not the value 0 (as confirmed by a low p-value) and the data relate linearly to the predictions (as indicated by a R^2^_adj_ close to 1). Where SEs were reported for BW and FEC (Höglund et al., 2018; O’Shaughnessy et al., 2015) and BW (Dimander et al., 2003), the deviations between data and prediction were generally in reasonable agreement with the estimated SEs (Fig. 6-8) except for the FEC in O’Shaughnessy et al. (2015); it is possible that the latter SEs are conservative indicators of uncertainty as they have constant value and may not fully account for overdispersion in egg counts. There was an exception, however, in the studies where the animals were inoculated with larger L3 doses (Dimander et al., 2003; Höglund et al., 2013); here, the magnitude of the peak of the FEC (at 5 weeks post turnout) was considerably underestimated by the model (Fig. 8), although there was agreement at the remaining time points; possible causes are discussed later.

**Table 6.**
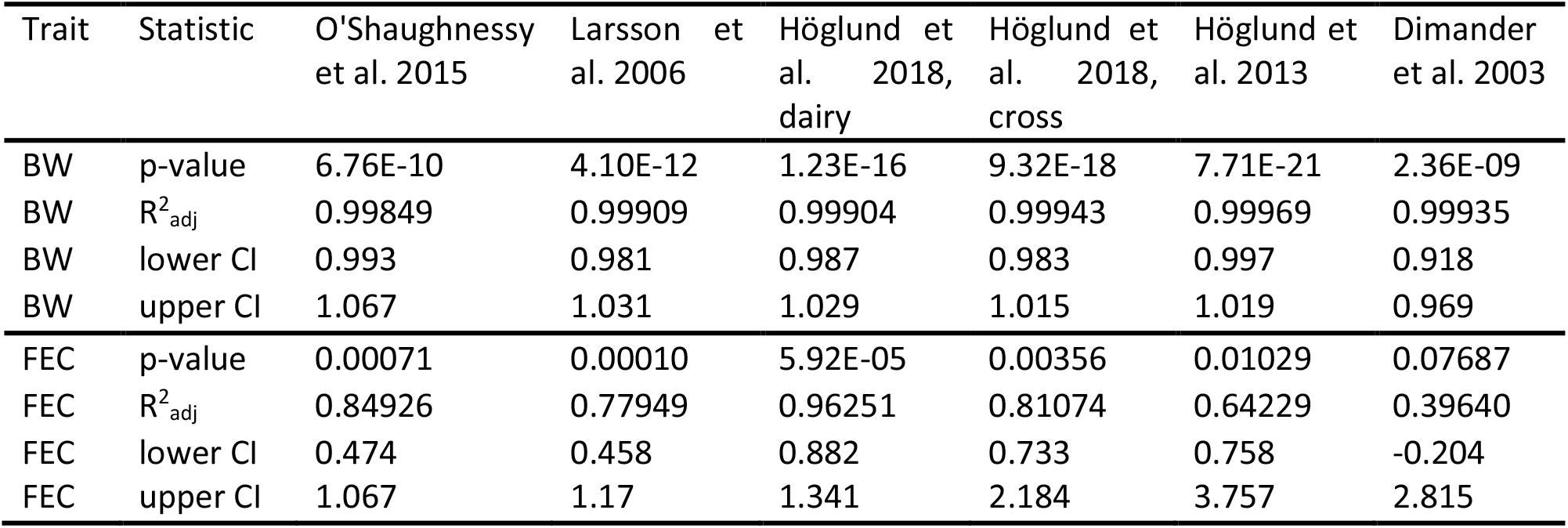
Statistical tests on the relatioship between the datasets and the model predictions. CI 95%: confidence interval of the slope of the linear relationship between observation data and prediction.

| Trait | Statistic | O'Shaughnessy et al. 2015 | Larsson et al. 2006 | Höglund et al. 2018, dairy | Höglund et al. 2018, cross | Höglund et al. 2013 | Dimander et al. 2003 |
| --- | --- | --- | --- | --- | --- | --- | --- |
| BW | p-value | 6.76E-10 | 4.10E-12 | 1.23E-16 | 9.32E-18 | 7.71E-21 | 2.36E-09 |
| BW | $R^2_{adj}$ | 0.99849 | 0.99909 | 0.99904 | 0.99943 | 0.99969 | 0.99935 |
| BW | lower CI | 0.993 | 0.981 | 0.987 | 0.983 | 0.997 | 0.918 |
| BW | upper CI | 1.067 | 1.031 | 1.029 | 1.015 | 1.019 | 0.969 |
| FEC | p-value | 0.00071 | 0.00010 | 5.92E-05 | 0.00356 | 0.01029 | 0.07687 |
| FEC | $R^2_{adj}$ | 0.84926 | 0.77949 | 0.96251 | 0.81074 | 0.64229 | 0.39640 |
| FEC | lower CI | 0.474 | 0.458 | 0.882 | 0.733 | 0.758 | -0.204 |
| FEC | upper CI | 1.067 | 1.17 | 1.341 | 2.184 | 3.757 | 2.815 |

#### 3.2.2 L3 on herbage

The predicted L3 concentration on herbage was in less quantitative agreement with the data (Fig. 6 and 8) (Dimander et al., 2003; Larsson et al., 2006; O’Shaughnessy et al., 2015) than the predicted BW and FEC. In two out of three cases where data were available, the model overestimated the magnitude of the observations considerably, although in all cases the predicted time trend seems consistent with that of the data. There are, however, many factors, some of which method-related, that can limit the efficacy and cause variability in the field recovery of L3 (Cain et al., 2021; Kloosterman, 1971; Paras et al., 2018; Tontini et al., 2019; Verschave et al., 2015). In particular, the fact one or two of the L3 counts suddenly dropped to zero and then rebounded during the season (Fig. 6) could result from sampling variation, although some of these sample points occurred when L3 availability was predicted by the model to have dropped suddenly and temporarily due to climatic factors. In O’Shaughnessy et al. (2015) (Fig. 6), the reported L3 refer to the SGS and not to the current season; in Dimander et al. (2003) (Fig. 8), L3 are reported for both the current and the following grazing seasons.

#### 3.2.3 Adult worms

The predictions of W were generally in reasonable agreement with the few data points from tracer animals (Fig. 6-8). In some cases (Table 5), these tracer animals grazed the same fields in a later grazing season, in which case the data may bear a weaker association to the original level of herbage contamination.

#### 3.2.4 Effective reproduction number

The patterns in R_e_ are more variable across the empirical studies than across the model behaviour examples due to the greater variation across studies, which includes differing weather, host growth and parasitological conditions. Yet, these patterns are similar among each other and to those in the behaviour examples, albeit being more variable in the extent and range of the oscillations above, near and occasionally below 1. Overall, the more or less precipitous drop in R_e_ through the grazing season is consistent with the challenge-dependent acquisition of immunity dampening the potential parasite population growth.

## 4. Discussion

We developed a novel mathematical model of the epidemiology of GIN infections in gazing animals. The model integrates variables describing the parasitic and FL stages of the nematode and the performance (growth, feed intake) and immunological states of the host. The model further includes compensatory host growth upon reduction in parasite load, the influence of weather on the parasite’s FL stage dynamics, and the influence of variable grass biomass on the ingestion of infectious parasite stages by the host. While the parasite population model uses a well-established framework for the dynamics of macroparasite infections (Anderson and May, 1978; Anderson and May, 1992) and follows previous attempts for GINs in cattle (Grenfell et al., 1987a; Roberts and Grenfell, 1991; Smith and Grenfell, 1994), the interactions with grass, weather data, and animal growth are novel and allow for the exploration of climate-driven effects and eventually the optimisation of treatment strategies based on performance as well as parasitological criteria. Therefore, we consider the inclusion of these variables central to the use and further development of models as tools to help address the challenges set out in the Introduction, i.e. the evaluation of alternative treatment and management strategies to current practices required by emerging anthelminthic resistance and climate change. Our first hypothesis, that the interactions within the model lead to interpretable nonlinear responses in the system that can be explored to enhance the outcomes of parasite control interventions, was supported by the study of model behaviour. Our second hypothesis, that the model is able to represent patterns of animal infection and performance in experimental trials, was supported by the outcomes of the model validation.

### 4.1 Modelling approach and scope

The model was parameterised, using literature sources, specifically for *O. ostertagi* in grazing cattle in temperate climates. We did so because of the clinical and economic importance of this species, particularly in young cattle (Armour, 1980; Charlier et al., 2020b; Forbes, 2020) and because of the greater knowledge of the relevant parameters for this species (Grenfell et al., 1987b; Michel, 1969; Michel et al., 1973; Rose et al., 2015; Smith et al., 1987; Verschave et al., 2014). In addition, model predictions were tested (see below) against datasets on FGS as this comprises all young cattle and in an attempt to develop and test the processes of acquisition of immunity from a known, naïve state, which avoids confounding effects from parasitological history.

We integrated, for the first time, processes relating to infection and immunity in cattle with the dynamics of the FL stages, grass availability and animal growth. Epidemiological models of the *O. ostertagi* lifecycle have been previously developed (for reviews of models of GINs in cattle see (Cornell, 2005; Smith and Grenfell, 1994; Verschave et al., 2016a)), but stopped short of incorporating all these factors. The first innovation added here is a model of the dynamics of parasite FL stages (Rose et al., 2015) that extended earlier work (Grenfell et al., 1986, 1987a; Smith, 1990; Smith et al., 1986) by including the influence of weather on soil-herbage migration, and to which we added egg shedding and larva ingestion by the cattle herd (sub-model 4). The second innovation is a model of the dynamics of the host state that builds on and adapts past work on parasite load and acquired immunity processes (Grenfell et al., 1987a, b; Roberts and Grenfell, 1991; Smith and Grenfell, 1994) (sub-model 1) and adds further variables describing host growth similarly to Vagenas et al. (2007) and Berk et al. (2016a); Berk et al. (2016b) (sub-model 2). Our host-state model differs from that of Berk in using fewer host state variables, distinct parameters and parameter values, and a revised representation of parasite-induced anorexia (Coop and Kyriazakis, 1999) on feed intake and the addition of compensatory growth. We note also that Berk’s model included a deliberately simplified representation of the parasite FL stages, whose availability varied seasonally but not according to egg output that developed under the influence of weather. Thirdly, our model includes (sub-model 3) dynamic variation in grass availability (Grenfell, 1988), which influences both animal growth and the concentration and hence the ingestion of parasite infective stages.

One novel aspect that emerged in this integrated model, is an explicit relationship of the rate of parasite transmission, or instantaneous rate of infection, Eq. (27), to variables relating to the host, parasite and grazing environment:

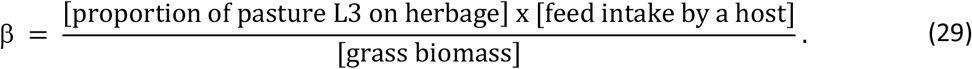

In Eq. (29), the proportion of L3 on herbage depends on the current weather; the feed intake depends on the current host body weight and parasite burden; and the grass biomass concentration depends on the grazing history and carrying capacity of the grazing system. Equation (29) builds on and adds to previous work (Anderson and May, 1978; Grenfell, 1988; Grenfell et al., 1987a; Kao et al., 2000; Louie et al., 2005; Singleton et al., 2011; Smith and Grenfell, 1985) a dynamic trade-off between host and environmental variables. Models are often very sensitive to the value of β when β is treated as a constant parameter (Grenfell et al., 1987a); in our model, however, β is a variable controlled by simultaneously-changing variables whose effects may either add or counterbalance. Based on the current parameters of the model, the variation of β during the grazing season was in the range 10^-3^ - 10^-4^/day/larva/host (c.f. model behaviour example in Supplementary Fig. S2), in agreement with estimates of β for *O. ostertagi* in cattle (Smith and Grenfell, 1985) and for other GINs in sheep (Kao et al., 2000).

As the processes modelled are not specific to *O. ostertagi* (including not being specific to its abomasal location, except possibly Eq. (23) the model has the potential to be re-parameterised for application to other parasites with a similar direct lifecycle, i.e. where transmission occurs through free-living eggs and larvae (Anderson and May, 1992; Smith and Grenfell, 1994). Such parasites include GINs in cattle such as *Cooperia* species. First steps have already been taken in extending the FL-stage dynamics (Grenfell et al., 1986; Sauermann and Leathwick, 2018) and the parasitic-stage dynamics (Rose Vineer et al., 2020b) to these species, although not yet in an integrated full-cycle model.

In principle, the model can also be adapted to GINs in other ruminant species. In fact, several full-cycle models have been developed for other ruminants (Verschave et al., 2016a) such as sheep (Kao et al., 2000; Louie et al., 2007; Singleton et al., 2011), incorporating similar essential mechanistic understanding and specific parameter estimates. Some of these models are simpler to analyse, parameterise and apply than the current model; however, most incorporate fewer state variables than would be necessary to describe host growth and immunity and the influence of weather and climate on parasite dynamics. The current model therefore offers advantages that might be extended to other systems, especially when seeking to predict outcomes and optimise interventions for both performance and parasite control, as recommended to attenuate the development of anthelmintic resistance (Charlier et al., 2014).

Given the many sources of uncertainty in the parasite and host dynamics, including uncertainty in the model parameters, reliable forecasting for a specific situation cannot be reasonably expected (Cornell, 2005; Grenfell et al., 1987a; Smith and Grenfell, 1994); this is even more so as strong influencers like weather and pasture contamination cannot be predicted at a future time. Instead, this and related models (Verschave et al., 2016a) are suited for predicting system responses to given parasite control strategies in order to classify their relative efficacies (Cornell, 2005; Smith, 2011). Therefore, agreement with observed patterns of infection and growth in published trials is important to build confidence in the use of the model under different conditions.

### 4.2 Model validation

We have demonstrated the ability of this new model to capture parasite and host dynamics in real systems through a validation exercise. Validation was carried out on empirical studies in Northern Europe reporting BW and FEC variables and containing a non-treated group. In addition, the animals in these studies were infected naturally through grazing such that the parasite dynamics were controlled by weather and host-parasite interactions alone. Some studies included inoculation of L3 at turnout, but subsequent infection was exclusively through grazing. The model predictions compared satisfactorily against the observations of BW and FEC across all studies, both graphically and in formal statistical testing. However, there were two studies, whose animals were inoculated with larger L3 doses at turnout, where the model underestimated considerably the magnitude of the FEC peak, although there was good agreement at the remaining time points. One explanation for this outcome stems from the fact that the animals were subjected to co-infection, predominantly by *O. ostertagi* and *C. oncophora* (Table 5), as is typical in natural field infections (Henriksen et al., 1976; Högberg et al., 2021; Michel et al., 1970), and that the FEC data used did not differentiate parasite species. As *C. oncophora* has considerably higher maximum fecundity (prior to being regulated) (Kloosterman et al., 1984; Verschave et al., 2016a; Verschave et al., 2014), a model parameterised for *O. ostertagi* would be expected to lead to lower FEC prediction, particularly in studies where parasite inoculated doses and loads are higher and at the peak of egg production, i.e. prior to the strong regulation of fecundity imposed by the developing acquired immunity and worm burden. A similar occurrence has been reported in previous model validation exercises (Smith and Guerrero, 1993). Likewise, experimental studies have suggested that FEC may differ between species at its peak but not necessarily at other time points (Hilderson et al., 1995; Kloosterman et al., 1984).

The use of studies with co-infection was imposed by a lack of data on single-species infections under natural weather conditions, which is required in order to test a full-cycle model. However, testing models under realistic field conditions, where co-infection by parasite species is common, can be regarded as desirable. Nevertheless, given that the model was designed and parameterised for a single parasite species, we, like others (Smith and Guerrero, 1993), are cautiously optimistic about the extent to which the model can represent co-infection situations. As we described earlier, the initial L3 concentrations input in the model and the parasite loads predicted were interpreted as representing total infection by both parasite species. Our working hypotheses were that: 1) there are no interactions between the species within the host, and, 2) the parameters and processes in the model are adequate for describing the dynamics of both species. Under these hypotheses, the model can be regarded as representing a typical parasite mixture, where e.g. *C. oncophora* dominates early and *O. ostertagi* dominates later (Dimander et al., 2003; Högberg et al., 2021). Regarding the first hypothesis, there is no experimental evidence of interaction between the host responses to *O. ostertagi* and *C. oncophora* in grazing calves (Dorny et al., 1997; Hilderson et al., 1995; Satrija and Nansen, 1993), although there is some evidence of cross-immunity (Kloosterman et al., 1984). There is evidence of interaction between co-infecting GINs of other species in cattle (Herlich, 1965) and in sheep and goats (Basripuzi et al., 2020; Lello et al., 2018; Sykes et al., 2009). Accounting for possible interaction between *O. ostertagi* and *C. oncophora* in cattle would require more experimental knowledge and further model advancement (see below).

Regarding the second hypothesis of adequacy of our model to describe both parasite species, it is likely that the same basic mechanisms are suitable to describe both species, at least at the level of simplification of the models, e.g. the location of establishment in the gastrointestinal tract, which differs between *O. ostertagi* and *C. oncophora*, is not specified in the model. However, there may be differences in parameter values between parasite species, e.g. in rate of acquisition of immunity (Dorny et al., 1997; Hilderson et al., 1995), although only some of the model parameters have been quantified for both species (Rose et al., 2015; Rose Vineer et al., 2020b; Verschave et al., 2016a; Verschave et al., 2014). One reason why the model may have approximated satisfactorily many of the variables in these studies, is that several of the parameters may be similar enough between the two parasite species (Grenfell et al., 1986; Rose Vineer et al., 2020b; Verschave et al., 2016b; Verschave et al., 2014), and some of those that differ more could have contrasting effects on the overall dynamics, e.g. through characteristics of the immune response vs fecundity, as suggested by experiments (Hilderson et al., 1995). Moreover, where there are differences, we expect them to be greater when egg production and parasite loads are higher, which, due to the regulatory effects of immunity and density-dependency, may be relatively short-lived and occur predominantly near the peak of FEC. Therefore, there is a cautious indication the model may, to a degree, be able to capture typical seasonally-varying mixtures of these parasites in the field, although this is an area where future research is clearly needed.

Compared to the predictions of BW and FEC, prediction of the L3 concentration on herbage was in less quantitative agreement with the data (where available); the model overestimated the abundance of L3 on pasture relative to that observed, although the patterns of the predicted time trends were consistent with those of the data. However, many factors can contribute to low efficiency and sampling variation in the field recovery of L3. These factors include the recovery method and the analyst (Cain et al., 2021; Kloosterman, 1971; Paras et al., 2018; Verschave et al., 2015), differing grass growth and under-sampling of the sward at the lowest level (Tontini et al., 2019), soil-herbage migration of L3, and avoidance of faecal pats or dung beetles, which associate with higher L3 concentration (Henriksen et al., 1976; Nansen et al., 1988). Measurement variation within a study can result from limited sampling of highly aggregated L3, and this possibility cannot be excluded in the studies where L3 counts dropped to zero and rebounded during the season. On the other hand, the use of differing recovery methods can lead to differences in recovery rate between studies (Verschave et al., 2015). Therefore, we would not regard the above overestimation as significant. The predictions of worm counts, W, were generally in agreement with the small number of post-mortem counts from tracer animals, some of which grazed the same fields concurrently while others did so in subsequent years.

A very small subset of model parameters or variables was informed by factors reported in the studies used for validation. Factors that were not measured in these experiments may have influenced the observations. These could include weather (beyond temperature and rainfall, which were included in the model); management; initial pasture contamination; immune status (naïve); faecal moisture content; sampling variation of the FEC method used; density and growth of the grass biomass; apparent digestibility of grass DM and use of feed supplements; and genetic strength and speed of the immune response. As we did not have information on any of these factors and we were not fitting the model to the data, we assumed that all remaining parameters of the model did not differ between studies. Given the potentially unaccounted-for variables, the ability of the model to produce estimates of parasite population and animal growth so close to observed values provides confidence in its ability to predict system dynamics under different, broader conditions.

### 4.3 Model behaviour

We have analysed some of the model behaviour by changing each of a small number of parameters. This analysis served to confirm expected qualitative outcomes and gain further confidence in the model, and to demonstrate some of the insights that can be derived from an integrated full-cycle model by exploring what-if scenarios. Some scenarios may be hypothetical or impractical to test experimentally in complex pasture systems, making the availability of models particularly valuable as analytical tools.

Changing the parasitological history of the pasture by increasing its contamination level at turnout led to earlier peaks in the predicted FEC and worm burden; these time shifts are similar to known peak-shift effects on the prevalence of macroparasites when increasing the force of infection on the host population (Anderson and May, 1985; Woolhouse, 1998). These results also agree with earlier model predictions (Berk et al., 2016b), except the latter contained two successive peaks, while we predicted a single peak during the season and none of the empirical FEC datasets used for validation indicated the occurrence of two peaks. The difference could stem from differing weather or from differing modelling of the parasite FL-stage dynamics, which in Berk et al. (2016b) excluded the influence of precipitation.

Likewise, changing in our model the size of the host population by increasing the cattle stocking density led to similar time shifts in the peak excretion of transmission stages and in the peak worm burden. The magnitude of the peak worm burden increased with increasing stocking density in agreement with earlier model predictions (Berk et al., 2016b; Grenfell et al., 1987a). We predicted this same pattern for the peak FEC, which agrees with Berk et al. (2016b) but is opposite to the pattern in Grenfell et al. (1987a), who highlighted that the worm burden, W, is a more indicative prediction as the FEC is known to be a poor index of parasite burden; although W is more rarely measured for obvious reasons.

Overall, these results confirmed expectations about the behaviour of the model; they also illustrate the importance of measuring L3 on pasture at turnout as some aspects of prediction can be uncertain if this variable is unknown. In addition, in each of the scenarios above there were appreciable effects on BW gain, with the differential between BW trajectories reducing by end of season (due to compensatory growth) in the case of differing initial contamination, but with little or no recovery in lost gain in the case of differing stocking densities. Likewise, the levels of L3 herbage contamination converged by the end of season in the first case, but diverged in the case of differing stocking densities. These outcomes are consistent with lasting effects of higher stocking density (Hansen et al., 1989; Thamsborg et al., 1998) and further confirm expected model behaviour. Overall, the above results highlight that the effects of parasitological history due to grazing in the previous season can be transient, while those of more intense grazing can dominate and be long lasting.

Exploring a set of simple drug treatment strategies, we obtained the following results. First, implementing a single anthelmintic treatment, the model predicted that the optimal application is at an intermediate time after turnout, e.g. 4 weeks, rather than immediately on turnout or later.in the season. This choice is based on multiple criteria: it led to the highest cumulative BW gain and to lower cumulative parasite burden and parasite excretion by the host and thus to potentially lower risk of clinical disease; on the other hand, herbage contamination was comparable to that in the late treatment. This outcome agrees with the expectation that delaying treatment to midseason allows the development of immunity and leads to better parasite control in the long run, while curbing the delay pre-empts the onset of parasite-induced anorexia and leads to better performance. Second, implementing treatment at turnout, the model predicted that the best timing for application of a second treatment is within a time window of 5 to 7 weeks after turnout, rather than immediately after the end of the first treatment at 3 weeks after turnout. This strategy led to a lower level of herbage contamination carried over to the next season and to lower cumulative parasite burden, although it did not lead to significant differences in performance. These treatment scenarios were illustrative and not chosen to mimic specific treatment regimens, although administration of persistent anthelmintic formulations early in the grazing season tend to be favoured due to their strong suppression of egg outputs and consequently of L3 levels. This strategy, however, has been posited to slow the acquisition of immunity (Vercruysse et al., 1994) and could therefore be counterproductive. Overall, these results illustrate the usefulness of a full-cycle epidemiological model for analysing and choosing treatment and management strategies, in particular accounting for performance and not only parasitological outcomes.

The predicted effective reproduction number, R_e_, exhibited a similar pattern across the empirical studies and the model behaviour analyses: very large initial values, a rapid decline due to the limiting effects of acquired immunity and density dependency, followed by narrow-ranged variation nearly containing the value of 1. We interpret this pattern as reassuring. First, across a range of differing parasitological, host and weather conditions, it agrees with the expectation that the parasite populations will have converged to a state of quasi stability superimposed by short-term fluctuations due to variable host response, weather and seasonal climate. Second, this result supports, rather than questions the consistency of the parameters of the host and free-living model components that we have attempted to integrate into a full-cycle model. Note that such consistency is not automatic as several parameters have been estimated independently.

### 4.4 Model assumptions and extensions

The model makes several simplifying assumptions already stated. One of the assumptions was that, to first approximation, the growth rate and DM content of the grass biomass did not vary with the weather and throughout the season. Such dependency could be included; however, a fuller account of environmental influence may involve further variables such as soil moisture saturation, and in turn soil type and topography, as well as management factors in relation to grass cultivar and fertiliser application. These refinements go beyond our current purpose and would require substantial empirical support. The current constant rate of grass growth is the average of empirical records from the location and period of the Baseline system (Table 3), but we expect variation in grass growth would have only a mild effect on the transmission rate β (Eq. (29)). The model also did not include the arrest or hypobiosis of parasitic larval stages and their subsequent re-emergence (Armour, 1980; Michel et al., 1976; Smith and Grenfell, 1985), although this would become relevant only towards the end of the FGS and beyond. Nevertheless, the higher late-season levels of L3 predicted under some scenarios could drive important epidemiological consequences, for example by causing higher risk of type II ostertagiosis through the re-emergence of arrested larvae, or by increasing the levels of pasture contamination in the following season through increased L3 emergence or L3 overwintering on pasture. The model could be extended to include hypobiosis, for example if applied to cattle in subsequent grazing seasons. The consequences of parasite exposure for immunity in older age classes could also be explored in an extended model, including the application of targeted treatment approaches in herds with differing levels of immunity (Ravinet et al., 2017). The model assumed the animals were under thermal neutrality by considering maintenance requirements that did not vary with temperature. The model can be extended to include thermal variation in intake, which is expected to be a very small fraction under varying moderate ambient temperature, but could increase in magnitude under climate warming scenarios.

The model describes the dynamics of an average animal and characterises the grazing population through its stocking density. In particular, the model does not include genetic and phenotypic variation. In fact, Smith and Guerrero (1993) have suggested that host heterogeneity in parasite load can be ignored in models aiming to evaluate control strategies that treat all animals in the same way. However, individual-based approaches have been evaluated for sheep (Louie et al., 2005) and cattle (Berk et al., 2016b). The current model could be extended to explore optimal strategies for targeted selected treatments based on individual infection or performance status (Charlier et al., 2014; Höglund et al., 2013; Merlin et al., 2017). Such host heterogeneity provides one of the mechanisms thought to generate the observed aggregation in parasite load and FEC among hosts, the other being the observed aggregation of L3 on pasture (Anderson and Gordon, 1982; Cornell et al., 2004). An alternative, empirical way of accounting for the latter is to make the number of ingested L3 a random variable with an empirical overdispersed distribution such as the negative binomial (Berk et al., 2016b; Smith and Guerrero, 1993) (we took this approach when comparing predictions with empirical data but not in the inherent parasite dynamics). Alternatively, individual-based formulations with stochastic dynamics and spatially-heterogeneous exposure allow both forms of aggregation to be linked mechanistically (Cornell, 2005; Cornell et al., 2004; Fox et al., 2013), but are usually applied to simpler representations of the GIN cycle for tractability (Smith and Grenfell, 1994), and the current addition of weather-driven variation will account for part of the system’s dynamic stochasticity. The inclusion of aggregation would strengthen the evaluation of control strategies further when it is relevant to account for heterogeneity in the parasite population; it is expected to influence the dynamics of invading anthelmintic-resistant strains and persisting non-resistant refugia (Cornell, 2005; van Wyk, 2001). Due to aggregation, and other factors already discussed, observations of L3 on herbage in the empirical studies (where reported) are uncertain; its potential effect could have been evaluated by generating distributions of predictions based on an assumed range of input values; these would be expected to include the data in the validation exercise. Sensitivity to uncertainty in other input parameters could be tackled similarly. Taking such an approach would have added an extra layer of complexity to the results, while sensitivity to such factors can be assessed already from the results of the model behaviour study.

Finally, as we already discussed extensively, the model was designed for infection by a single-parasite, i.e. *O. ostertagi*, although it was applied to co-infections for reasons explained. This is the case of most models developed for specific GIN infections. Our results, however, supported the application to co-infections by *O. ostertagi* and *C. oncophora* in the empirical studies considered here. A future challenge is to extend such nonlinear models to account explicitly for co-infection. Generic models investigating the implications of parasite co-infection have, for tractability, assumed unspecific host responses to parasite burdens and thus that parasite species did not interact directly (Dobson and Roberts, 1994), but there have been theoretical attempts at including such effects (Bottomley et al., 2005). However, currently, there is little knowledge about which responses would be interacting and how; therefore, more empirical study on GIN co-infection is needed.

### 4.5 Conclusions

We developed a model of the full life-cycle of *O. ostertagi* that for the first time also incorporates grass and animal growth and data-driven environmental effects on infective larval availability, in addition to host immunity. The model was able to reproduce expected patterns and scales of host growth and parasite dynamics in first season grazing cattle, and closely matched observed results in published studies without the need for model fitting. Exploration of initial pasture conditions, stocking density and treatment scenarios showed that the model can be used to predict the effects of management and climate on infection patterns. Future application could include optimisation of intervention strategies under rapidly changing climate and advancing anthelmintic resistance.

## Acknowledgements

This research was supported by funding from UK Research and Innovation through grant BB/R010250/1. JANF was also supported in part by the Scottish Government’s Rural and Environment Science and Analytical Services (RESAS). The funders had no role in study design, model development, data collection and analysis, decision to publish, or preparation of the manuscript.

## Declaration of interest

None to declare.

## CRediT authorship contribution statement

**Joao A. N. Filipe:** Conceptualisation, Funding acquisition, Data curation, Formal analysis, Methodology, Project Administration, Software, Validation, Visualisation, Writing – original draft, Writing - review & editing. **Ilias Kyriazakis:** Conceptualisation, Funding acquisition, Project Administration, Validation, Writing – review & editing. **Christopher McFarland:** Investigation, Validation, Writing - review & editing. **Eric R. Morgan:** Conceptualisation, Funding acquisition, Project Administration, Resources, Validation, Writing - review & editing.

## Research data for this article

Data that were used to perform the study are publicly available or stated within the main text and in the Supplementary data. Code for the model is available (Filipe, 2022).

## Appendix A. Supplementary data

Supplementary data to this article can be found online at xxx

## Supplementary data 1

Text S1: Derivation of new model parameters: Cost of maintenance, Cmaint

Text S2: Derivation of new model parameters: Cost of acquired immunity resources, C_I1_ and C_I2_

Text S3: Gastrointestinal tract capacity

Text S4: Body weight drop at turnout

## Supplementary data 2

Table S1: Parasite free-living stages: environmental dependency

Table S2: Parasite free-living stages: initial state

## Supplementary data 3

Figure S1: Weather data used in all studies

Figure S2: Rate of transmission β

## Notes

### Competing Interest Statement

The authors have declared no competing interest.

