## Supplementary Material for "Novel epidemiological model of gastrointestinal-nematode infection to assess grazing cattle resilience by integrating host growth, parasite, grass and environmental dynamics"

### Supplementary Text

#### List of Contents

**Supplementary Text S1: Derivation of new model parameters: Cost of maintenance, $C_{maint}$**

**Supplementary Text S2: Derivation of new model parameters: Cost of acquired immunity resources, $C_{I1}$ and $C_{I2}$**

**Supplementary Text S3: Gastrointestinal tract capacity**

**Supplementary Text S4: Body weight drop at turnout**

##### Supplementary Text S1: Derivation of new model parameters: Cost of maintenance, $C_{maint}$

An estimate of parameter $C_{maint}$ in the rate of biomass use for maintenance functions ($D_{maint} = C_{maint}\ BW^{0.75}$, Eq. (16), Table 2) was obtained as follows, without evoking energy requirements explicitly. Work on genetic selection in cattle (Koch et al., 1963; Archer et al., 1997) has used the lowest residual feed intake (RFI), i.e. the difference between observed and predicted individual intake, as a selection criterion. The predicted average daily DM feed intake (ADFI) of an individual is often a linear model of the individual's average daily BW gain (ADG), the average daily DM intake required for maintenance (proportional to $BW^{0.75}$), and possibly other covariates, taking the form: $ADFI = a + b_1\ BW^{0.75} + b_2\ ADG + \ldots$, where a, $b_1$, $b_2$ are regression coefficients. The quantity $(b_1/b_2)\ BW^{0.75}$ gives an estimate of the cost of maintenance on a scale comparable to ADG. In addition, in line with Eq. (14), we accounted for the fact ADG is wet gain by multiplying by the average of $p_{dry}$, which over the range of BW in the current model is around 0.75. Literature estimates of $b_1$ and $b_2$ (Tedeschi, 2006; Cruz et al., 2010; Old et al., 2015) led to estimates 0.036, 0.026, and 0.020 for $b_1/b_2$, but the more recent analysis across published datasets (Old et al., 2015) gives extra confidence to the first estimate. Multiplying by 0.75, gave $C_{maint} = 0.03\ kg^{0.25}$ DM/d (Table 2). Our exploration showed that model behaviour was qualitatively suitable for $C_{maint}$ based on the above values of $b_1/b_2$, but BW and FI became unrealistic for lower values, offering some degree of consistency between the model and these estimates.

##### Supplementary Text S2: Derivation of new model parameters: Cost of acquired immunity resources, $C_{I1}$ and $C_{I2}$

An estimate of parameter $C_{I2}$ in the rate of biomass use for maintaining the acquired level of immunity ($C_{I2}\ I_m$ Eq. (17), Table 2) was derived from published measurements of the metabolisable protein requirement for the expression of immunity to *Teladorsagia circumcincta* in sheep (Houdijk et al., 2001). We assumed that these requirements can be transposed to *O. ostertagi* in cattle. The measured requirement of protein, $\Delta P$, is 0.7 g/kg $BW^{0.75}$/d. Assuming that there is a corresponding gain in body water ($\Delta W$), we quantified $\Delta W$ by differentiating allometric relationships between W and BW or P and BW as in Eq. (18) (Filipe et al., 2018), giving: $\Delta W = \rho\ \Delta P$, where $\rho = (b_W/b_P)\ (a_W/a_P)\ BW^{bW-bP}$, and $a_P$=1.697, $b_P$=0.601 and $a_W$=1.997, $b_W$=0.707 are estimates of these allometric parameters for cattle (Carstens et al., 1991) (Table 2). The total daily biomass intake for maintaining a maximum level of immunity is therefore $(\Delta W+\Delta P)\ BW^{0.75} = (\rho+1)\ \Delta P\ BW^{0.75}$. Using the values of the allometric parameters and assuming a BW of 700kg, gives $C_{I2}$ = 0.359, which we rounded to $C_{I2}$ = 0.4 kg/uI/d (Table 2).

For the rate $C_{I1}$ of biomass use per increase $dI_m/dt$ in the immunity level ($C_{I1}\, dI_m/dt$, Eq. (17)), we made the working assumption that the rate of biomass use for increase in $I_m$ is proportional to that for maintenance of $I_m$, i.e. $C_{I1}\, dI_m/dt = \varepsilon\, C_{I2}\, I_m$, giving $C_{I1} = \varepsilon\, C_{I2}\, I_m/(dI_m/dt)$. At half of the maximum level of immunity, i.e. $I_m$=0.5, an estimate of $dI_m/dt$ with the current values of the model parameters is 0.000877 uI/d, which leads to $C_{I1} = \varepsilon\ 57.0\ C_{I2}$. Assuming that $\varepsilon$=0.5, i.e. that it costs more to maintain than to mount immunity, we obtain $C_{I1}$ = 11.4, which we rounded to $C_{I1}$ = 10 kg/uI (Table 2). Here, uI is a unit of immunity, which corresponds to $I_m$ =1 in the current model.

##### Supplementary Text S3: Gastrointestinal tract capacity

The daily feed intake of a grazing animal was constrained by the capacity of the gastrointestinal tract. This capacity was represented by an almost linear relationship to BW: $G_{cap} = a_{cap}\ BW^{cap}$, where $a_{cap} = 10^{-0.936}$, $b_{cap}$ = 1.032 (Demment and Van Soest, 1985; Clauss et al., 2007). The actual feed intake DM was:

$$\mathrm{FIDM_{actual}} = \ \mathrm{minimum}\left(\mathrm{G_{cap}}, \frac{\mathrm{A_{out}FIDM}}{\mathrm{p_{dry}}}\right)\mathrm{p_{dry}}, \qquad \text{(A.1)}$$

where FIDM is given by Eq. (20) and $A_{out}$, if different from 1, is given in Text S4.

##### Supplementary Text S4: Body weight drop at turnout

A rapid and temporary drop in FI due to adaptation to grazing at turnout, i.e. movement from housing onto pasture at the start of the grazing season (Balch and Line, 1957; Fox et al., 1989), was represented by the function of time t since turnout:

$$\mathrm{A_{out}} = \exp(\ \log(\mathrm{n_{out}})\ \exp(-\mathrm{k_{out}}\ t + \mathrm{a_{out}})\ \left(\frac{\mathrm{k_{out}}\ t}{\mathrm{a_{out}}}\right)^{\mathrm{a_{out}}}. \qquad \text{(A.2)}$$

This function has a start value of 1, followed by a sharp drop (controlled by $a_{out}$) to a minimum value of $n_{out}$, and then bounces back to the value of 1 at rate controlled by $k_{out}$. The function was included as an additional multiplication factor in Eqs. (13), (14) and (20) in the cases listed at the end of Section 2.3. In the baseline system used to study model behaviour, $n_{out}$ = 0.5, $k_{out}$ = 1/3.04 (Balch and Line, 1957), and $a_{out}$ = 0.01, which causes an almost instantaneous drop to $n_{out}$. In other cases, the values were determined by BW observations.

### Supplementary Tables

**Supplementary Table S1. Parasite free-living stages.** Environmental dependency of the lifecycle parameters (Table 4) of the FL stage model (Rose et al., 2015). T is daily average temperature (°C) and P is daily precipitation (mm/d). We replaced $\delta$ for $2*\delta$ in the original formulation.

| Parameter | Description |
|---|---|
| $\delta$ | $2*(- 0.07258 + 0.00976*T)$ |
| $\mu_1$ | $\exp(-4.38278 - 0.1064*T + 0.0054*T^2)$ |
| $\mu_2$ | $\mu_1$ |
| $\mu_1$ | $10\ \mu_4$ |
| $\mu_4$ | $\exp(-6.388 - 0.26810*T + 0.01633*T^2 - 0.00016*T^3)$ |
| $\mu_5$ | $\mu_3$ |
| $m_1$ | 0 if P<2, 0.06 otherwise |
| $m_2$ | $\exp(-5.4824 + 0.45392*T - 0.01252*T^2)$ |

**Supplementary Table S2. Parasite free-living stages.** Initial values of the parasite's state variables (Table 4) at turnout (t=0).

| Variable | Value | Units | Comments |
|---|---|---|---|
| $E_p$ | 0 | eggs/ha | Assuming no overwinter survival |
| $E_c$ | 0 | eggs/ha | idem |
| $L_{12}$ | 0 | larvae/ha | idem |
| $L_{3f}$ | 0 | larvae/ha | idem |
| $L_{3p}$ | $L_{3c}(0)\ G(0)/m_2(0)$ | larvae/ha | Assuming initial contamination $L_{3c}(0)$ |

### Supplementary Figures

#### List of Contents

**Supplementary Figure S1: Weather data used in all model predictions**

**Supplementary Figure S2: Parasite transmission rate β during the grazing season**

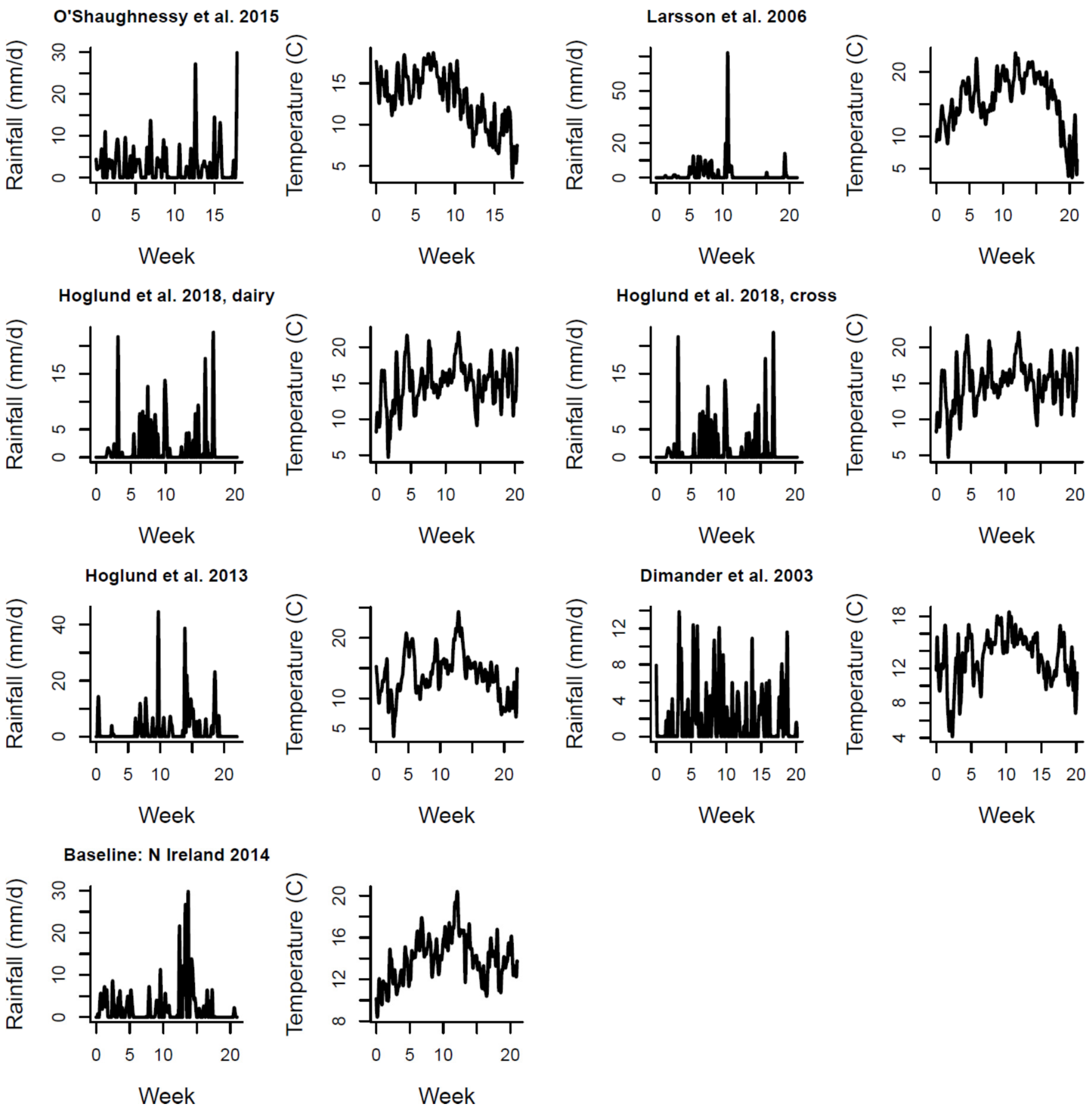


**Fig. S1. Weather variables used in all model predictions.** Daily average temperature (°C) and daily rainfall (mm/d) during the grazing periods of the six empirical studies used for model validation (Fig. 1-3) and of the baseline system (AFBI Hillsborough, Northern Ireland) used for the study of model behaviour (Fig. 4-7). The geographic locations and source of data are detailed in Sec. 8 and 9. Note axis scales differ between locations.

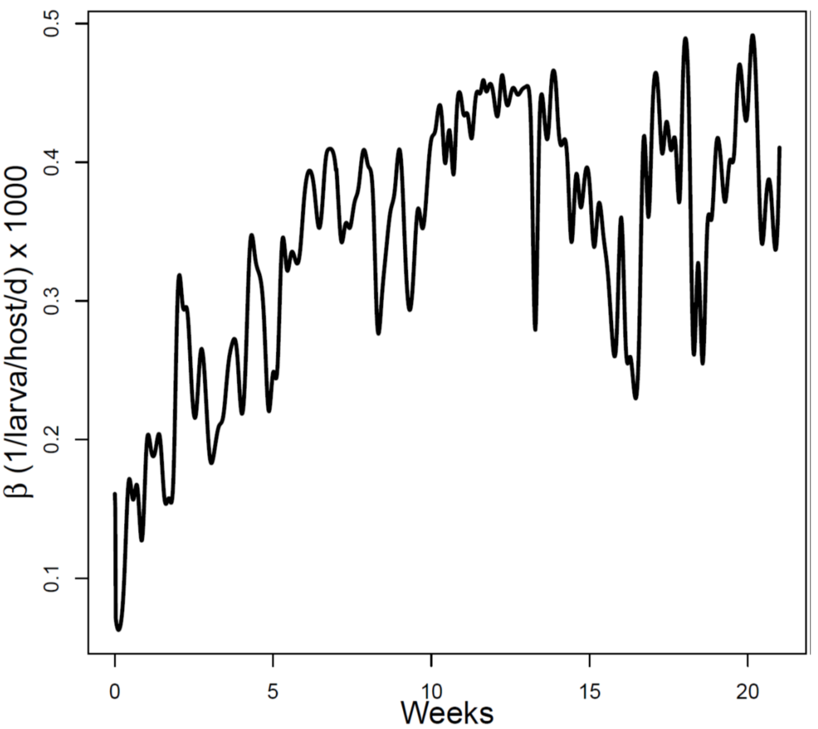


**Fig. S2. Parasite transmission rate β during the grazing season.** Transmission rate calculated as in Eq. (26) and (29). Case of model behaviour (baseline weather, Fig. A1) with two rounds of anthelmintic treatment applied at turnout and 7 week later (Sections 2.7.3 and 3.1.4Sec). The order of magnitude agrees with literature values (Section 4.1). In the other behaviour scenarios (Section 3) β has a similar temporal pattern and a magnitude that was no more than 20% above or below the current magnitude.
